# The αC-β4 loop controls the allosteric cooperativity between nucleotide and substrate in the catalytic subunit of protein kinase A

**DOI:** 10.1101/2023.09.12.557419

**Authors:** Cristina Olivieri, Yingjie Wang, Caitlin Walker, Manu V. Subrahmanian, Kim N. Ha, David A. Bernlohr, Jiali Gao, Carlo Camilloni, Michele Vendruscolo, Susan S. Taylor, Gianluigi Veglia

## Abstract

Allosteric cooperativity between ATP and substrates is a prominent characteristic of the cAMP-dependent catalytic subunit of protein kinase A (PKA). Not only this long-range synergistic action is involved in substrate recognition and fidelity, but it is also likely to regulate PKA association with regulatory subunits and other binding partners. To date, a complete understanding of the molecular determinants for this intramolecular mechanism is still lacking.

Here, we integrated NMR-restrained molecular dynamics simulations and a Markov State Model to characterize the free energy landscape and conformational transitions of the catalytic subunit of protein kinase A (PKA-C). We found that the apoenzyme populates a broad free energy basin featuring a conformational ensemble of the active state of PKA-C (ground state) and other basins with lower populations (excited states). The first excited state corresponds to a previously characterized inactive state of PKA-C with the αC helix swinging outward. The second excited state displays a disrupted hydrophobic packing around the regulatory (R) spine, with a flipped configuration of the F100 and F102 residues at the αC-β4 loop. To experimentally validate the second excited state, we mutated F100 into alanine (F100A) and used NMR spectroscopy to characterize the structural response of the kinase to ATP and substrate binding. While the catalytic efficiency of PKA-C^F100A^ with a canonical peptide substrate remains unaltered, this mutation rearranges the αC-β4 loop conformation, interrupting the structural coupling of the two lobes and abolishing the allosteric binding cooperativity of the enzyme. The highly conserved αC-β4 loop emerges as a pivotal element able to control the synergistic binding between nucleotide and substrate. These results may explain how mutations or insertions near or within this motif affect the function and drug sensitivity in other homologous kinases.

## INTRODUCTION

Eukaryotic protein kinases (EPKs) are plastic enzymes of paramount importance in signaling processes, catalyzing phosphoryl transfer reactions, or acting as scaffolds for other enzymes and/or binding partners ^1,2^. Of all kinases, the catalytic subunit of PKA (PKA-C) was the first to be structurally characterized by X-ray crystallography.^3,4^ In its inhibited state, PKA-C assembles into a heterotetrametric holoenzyme comprising two catalytic (C) and two regulatory (R) subunits.^5^ The canonical activation mechanism of PKA involves the binding of two cAMP molecules that disassemble the holoenzyme, unleashing active PKA-C monomers, which target signaling partners.^6^ In 2017, however, Scott and coworkers proposed an alternative activation mechanism in which the holoenzyme does not disassemble under physiological conditions; rather, it is clustered in *signaling islands*, localized near the substrates by A-kinase anchoring proteins (AKAPs), facilitating targeted signaling interactions.^7^ To date, the activation mechanism of PKA is still under active investigation since PKA-C appears to be fully liberated under pathological conditions.^8–10^

X-ray crystallography studies revealed that PKA-C is a bilobal enzyme, with a dynamic N-terminal lobe (N-lobe) comprising four β-sheets and an αC helix, while the C-terminal lobe (C-lobe) is more rigid and mostly composed of α-helices (**Figure 1A**).^3,4,11^ The N-lobe harbors the nucleotide-binding site, whereas the substrate binding cleft lays at the interface between the N- and C-lobe. The three-dimensional structure of PKA-C features a highly conserved hydrophobic core decorated with catalytically important motifs, *i.e.*, the Gly-rich loop, DFG-loop, activation loop, and magnesium and peptide positioning loops.^12^ In the catalytically active state, these motifs are all poised for phosphoryl transfer,^13–15^ a condition necessary but not sufficient to define an active kinase. More recent studies revealed a critical role of the enzyme’s hydrophobic core, which is crossed by a catalytic (C) spine and a regulatory (R) spine surrounded by shell residues (**Figure 1B**).^16,17^ The C spine comprises an array of hydrophobic residues and is assembled upon binding ATP, whereas the R spine is engaged when the activation loop is phosphorylated at Thr197, which positions the αC-helix in an active configuration.^18^

**Figure 1.**
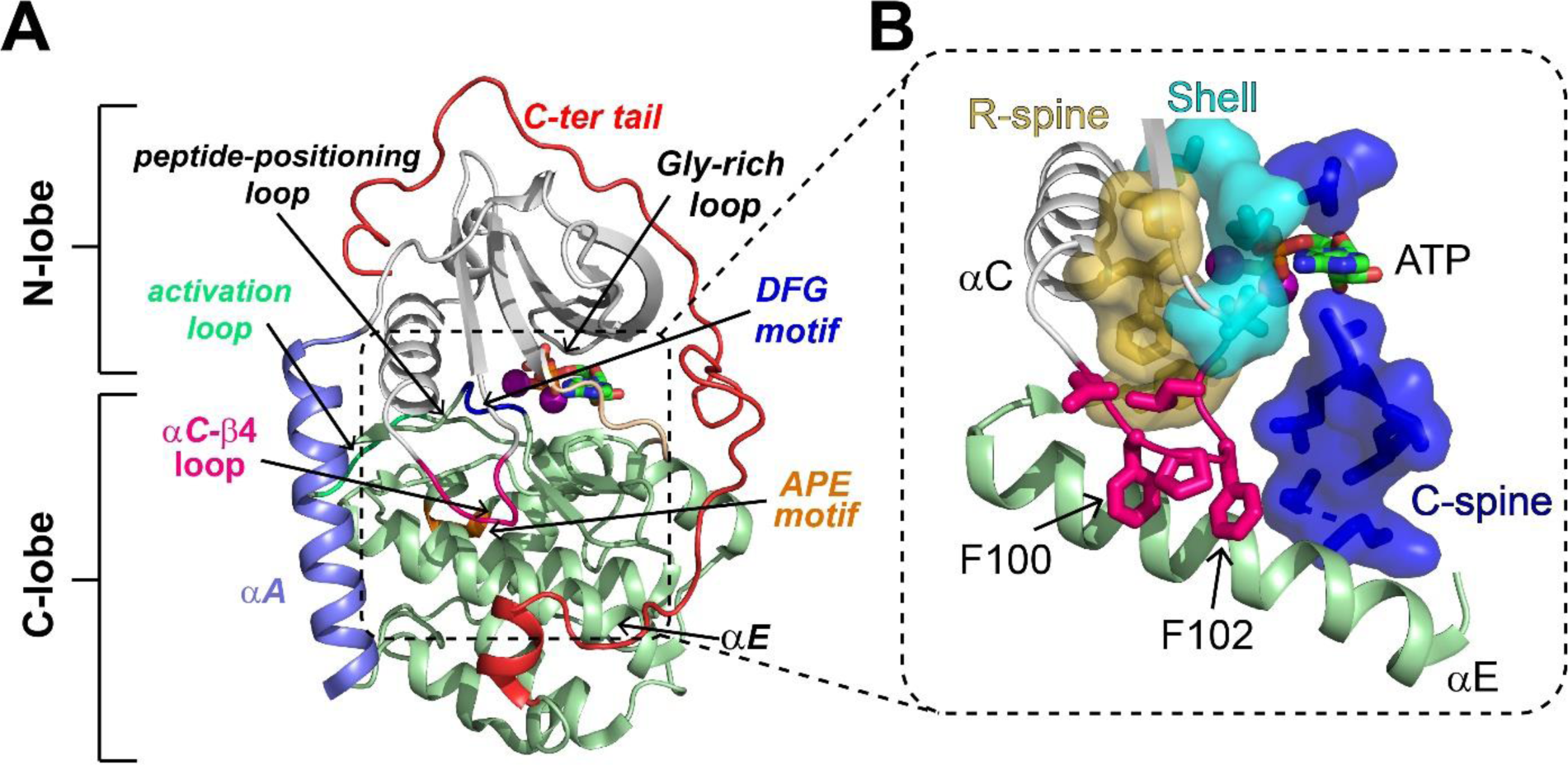
Structural and catalytic motifs of PKA-C. (**A**) Backbone representation of the ternary complex of PKA-C bound to ATP and the PKI_5-24_ peptide (not depicted), PDB Code 4WB5. Highlighted are key motifs including the αA, αC, and αE helices, C-terminal tail, activation loop, peptide-positioning loop, Gly-rich loop, and the DFG and APE motifs. (**B**) The hydrophobic core of PKA-C features the regulatory spine (R-spine, gold), catalytic spine (C-spine, blue), shell residues (cyan), and the αC-β4 loop (hot pink), which locks the αE helix and couples the two lobes of PKA-C.

A distinct property of PKA-C is the binding cooperativity between ATP and substrate.^19^ In the catalytic cycle, the kinase binds ATP and unphosphorylated substrates with positive binding cooperativity, whereas a negative binding cooperativity between ADP and phosphorylated substrate characterizes the exit complex.^20^ The biological importance of these phenomena has been emphasized by our recent studies on disease-driven mutations,^21–23^ which all feature disrupted cooperativity between nucleotides and protein kinase inhibitor (PKI) or canonical substrates.^21–23^ Since the recognition sequence of substrates is highly homologous to that of the R subunits,^5^ a loss of cooperativity may affect not only substrate binding fidelity but also regulatory processes. Thus far, the molecular determinants for the binding cooperativity between nucleotide and sub-strate and its role in PKA signalosome remain elusive.

Aiming to define the allosteric mechanism for PKA-C cooperativity, we combined NMR-restrained replica-averaged metadynamics (RAM)^24,25^ and Markov State Model (MSM)^26^ and charted the conformational landscape and dynamic transitions of the main forms of PKA-C. We found that the apo kinase occupies three distinct basins: (*i*) a most populated ground state (GS) with constitutively active conformations competent for catalysis, (*ii*) a first high free energy basin representative of typical inactive states with a dislodged configuration of the αC helix (ES1), and (*iii*) a second high free energy basin with a disrupted hydrophobic array of residues at the core of the enzyme (ES2). Notably, the equilibrium between the most populated ground state and the other low-populated states agrees with previous NMR Carr-Purcell-Meiboom-Gill (CPMG) relaxation dispersion and chemical exchange saturation transfer (CEST) measurements.^27^ To compound the existence of the second inactive basin, we mutated F100 at the αC-β4 loop into Ala (PKA-C^F100A^) and characterized its ligand binding thermodynamics and structural response by isothermal titration calorimetry (ITC) and NMR spectroscopy, respectively. We found that PKA-C^F100A^ phosphorylates a canonical peptide with a catalytic efficiency similar to the wild-type kinase (PKA-C^WT^); however, this mutation abolishes the binding cooperativity between nucleotide and substrate. The NMR experiments reveal that F100A perturbs the hydrophobic packing around the αC-β4 loop and interrupts the allosteric communication between the two lobes of the enzyme. Overall, these results emphasize the pivotal role of the αC-β4 loop in kinase function and may explain why single-site mutations or insertion mutations stabilizing this motif in homologous kinases result in oncogenes and confer differential drug sensitivity.^28–30^

## RESULTS

### The free energy landscape of PKA-C charted by NMR-restrained replica-averaged metadynamics (RAM) simulations

To characterize the experimentally accessible conformational landscape of PKA-C in the µs-to-ms timescale, we performed NMR-restrained metadynamics simulations within the RAM framework.^24,25^ Using four replicas restrained with backbone chemical shifts (CS), we simulated the apo, binary (PKA-C/ATP), and ternary (PKA-C/ATP/PKI_5-24_) complexes of PKA-C. Incorporating the CS restrains in the simulations ensures a close agreement with the conformational space explored by the kinase under our NMR experimental conditions,^31,32^ whereas the enhanced sampling through metadynamics simulations boosts the conformational plasticity of the enzyme along different degrees of freedom (*i.e.*, collective variables, CVs. See Methods) (**Figure 2 - figure supplement 1**).^33^ In our case, back-calculations of the CS values with Sparta+ ^34^ show that the restrained simulations improved the agreement between theoretical and experimental CS values of ∼0.2 ppm for the amide N atoms and ∼0.1 ppm for the remainder backbone atoms (**Figure 2 - figure supplement 2**). Also, the bias-exchange metadynamics allow each replica to span a significantly broader conformational space relative to classical MD simulations (**Figure 2 - figure supplement 3**). The metadynamics simulations were equilibrated and converged in 300 ns (**Figure 2A**, **Figure 2 - figure supplement 4**). The full free energy landscape was then reconstructed by sampling an extra 100 ns of production phase with reduced bias along each CV. We found that the population of the PKA-C conformers is modulated by ATP and pseudosubstrate binding within the NMR detection limit of sparsely populated states (∼ 0.5% or ΔG < 3.2 kcal/mol).

**Figure 2.**
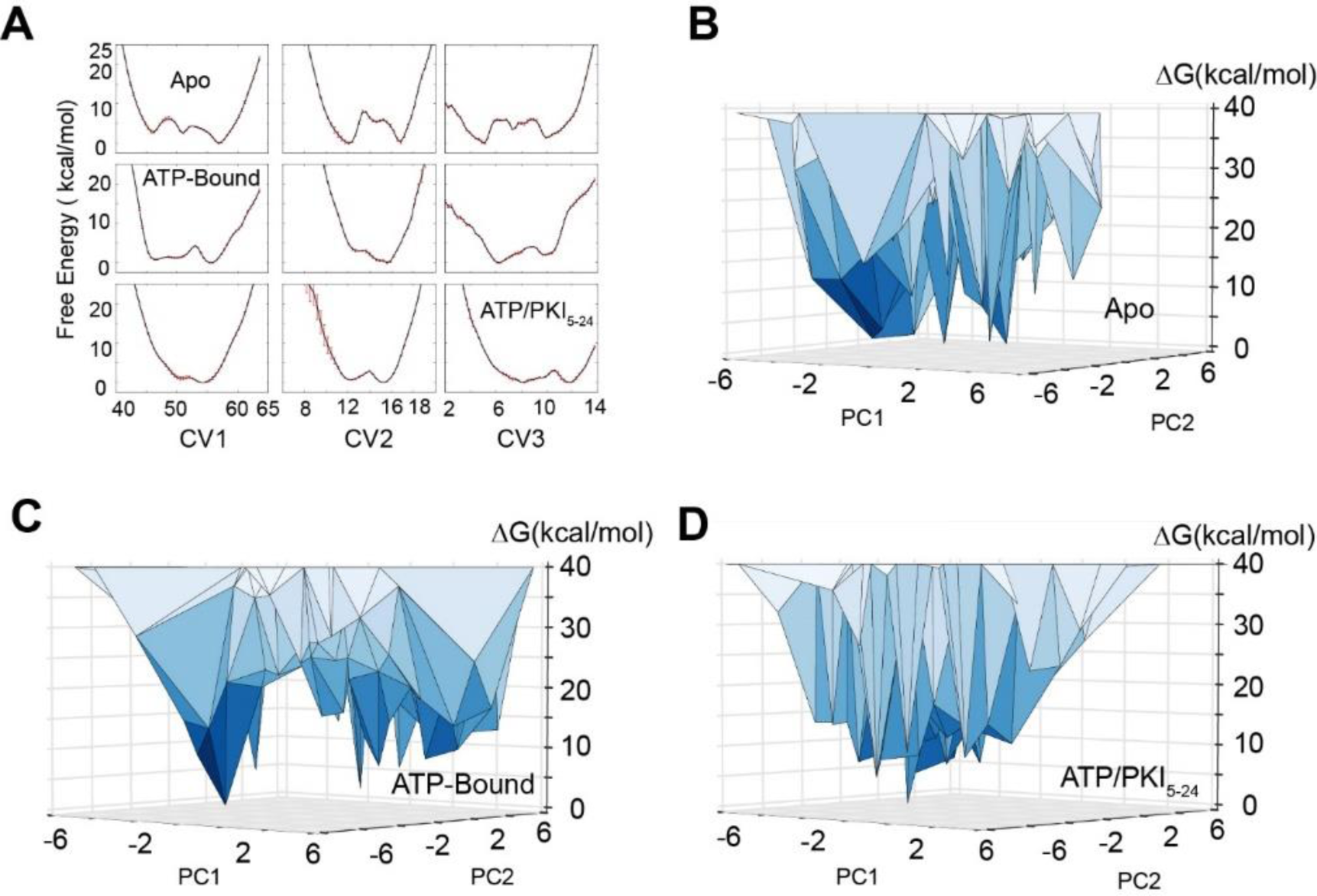
Free energy landscape of PKA-C obtained from replica-averaged metadynamics (RAM) simulations. (**A**) Convergence of the bias deposition along the first three collective variables (CVs). The free energy (expressed in kcal/mol) of the different CVs was averaged over the last 100 ns of RAM simulations. The standard deviations are reported as red error bars. (**B-D**) Free energy landscape along the first two principal components (PC1 and PC2) of PKA-C in the apo, ATP-bound, and ATP/PKI bound forms. PC1 and PC2 are projected from the first three CVs. The vertices represent conformational states. In the apo form, multiple states have comparable free energy with ΔG < 5 kcal/mol, whereas in the binary form, fewer states have ΔG < 5 kcal/mol. For the ternary form only a major ground state is populated.

Specifically, the apo PKA-C populates preferentially a ground state and five readily accessible low-populated excited states (**Figure 2B**, **Figure 2 – supplementary table 1**). The binary complex populates the same ground state and a broad higher energy basin (**Figure 2C**, **Figure 2 – supplementary table 1**). Finally, the ternary complex occupies a narrower dominant ground state (**Figure 2D**, **Figure 2 – supplementary table 1**). Overall, these calculations revealed that the apo kinase explores multiple minima along each CV. The binding of ATP shifts the population of the ensemble into an additional minimum, featuring structures that are committed to substrate binding. Finally, the conformational heterogeneity of the ensemble is significantly reduced upon formation of the ternary complex, which represents a catalytically committed state^13^ (**Figure 2A**). The free energy landscape obtained from these RAM simulations is consistent with the qualitative picture previously inferred from our NMR spin relaxation experiments and hydrogen/deuterium exchange studies,^35–37^ while providing a detailed structural characterization of the ground and sparsely populated conformationally excited states.

### MSM reveals the conformational transitions of PKA-C from ground to high free energy (excited) states

To refine the free energy landscape and delineate the kinetics of conformational transitions of the kinase, we performed additional unbiased sampling to build Markov State Models (MSM).^38,39^ MSMs are commonly used to describe the dynamic transitions of various metastable states for macromolecules in terms of their populations and transition rates. MSMs are typically created by combining thousands of short unbiased simulations.^38,39^ Following this strategy, we performed several short simulations (10 – 20 ns) using thousands of the low free energy conformations (ΔG < 3.2 kcal/mol) chosen randomly from the three forms of PKA-C as starting structures. The conformational ensembles were clustered into microstates and seeded to start a second round of adaptive sampling (see Methods). This iterative process was repeated three times to assure convergence and yielded 100 μs trajectories for both the apo and binary forms, whereas 60 μs trajectories were collected for the less dynamic ternary complex. Once we reached a sufficient sampling, we built an MSM including L95, V104, L106, M118, M120, Y164, and F185 to investigate the dynamic transitions of the hydrophobic R spine and shell residues (**Figure 3 – figure supplement 1**). These residues are ideal reporters of the dynamic processes governing the activation/deactivation of the kinase.^40^ To compare the free energy landscape of different complexes, we projected the conformational ensembles of three forms and existing crystal structures of PKA-C onto the first two time-lagged independent components (tICs) of the apo form, which were obtained by a time-lagged independent component analysis (tICA). These tICs represent the directions of the slowest motion of the kinase and visualize the conformational transitions of V104, L95, and F185 (**Figure 3 – figure supplement 1**). The MSM shows that the hydrophobic core of the apo PKA-C accesses three major basins. The broadest basin represents the ground state (GS) and corresponds to the conformations of PKA-C captured in several crystal structures (**Figure 3A**). The other two states (conformationally excited states) are less populated. The first excited state (ES1) features a disrupted hydrophobic packing of L95, V104, and F185. This conformation matches an inactive state with an outward orientation of the αC-helix typical of the inhibited states of PKA-C (**Figure 3 – Figure supplement 2**). The second state (ES2) displays a flipped configuration of the V104 side chain and disrupted hydrophobic interactions responsible for anchoring the αC-β4 loop to the C-lobe, with F100 and F102 adopting a gauche+ configuration and forming a stable π-π stacking. (**Figure 3A**). In contrast, the active GS ensemble features a *trans* configuration for the F100 and F102 side chains that stabilize the π-π stacking with Y156 at the αE-helix and cation-π interactions with R308 at the C-tail (**Figure 3A**). To our knowledge, the ES2 state was never observed in the available crystal structures. Upon ATP binding, the conformational space spanned by the kinase becomes narrower, and the conformers populate mostly the GS, with a small fraction in the ES1 state (**Figure 3B**). This is consistent with the role of the nucleotide as an allosteric effector that enhances the affinity of the enzyme for the substrate.^41^ In the ternary form (ATP and PKI-bound), PKA-C populates only the GS consistent with the competent conformation observed in the ternary structure (*i.e.*, 1ATP). In this case, the αC-β4 loop of the enzyme is locked in a well-defined, active configuration (**Figure 3A**).

**Figure 3.**
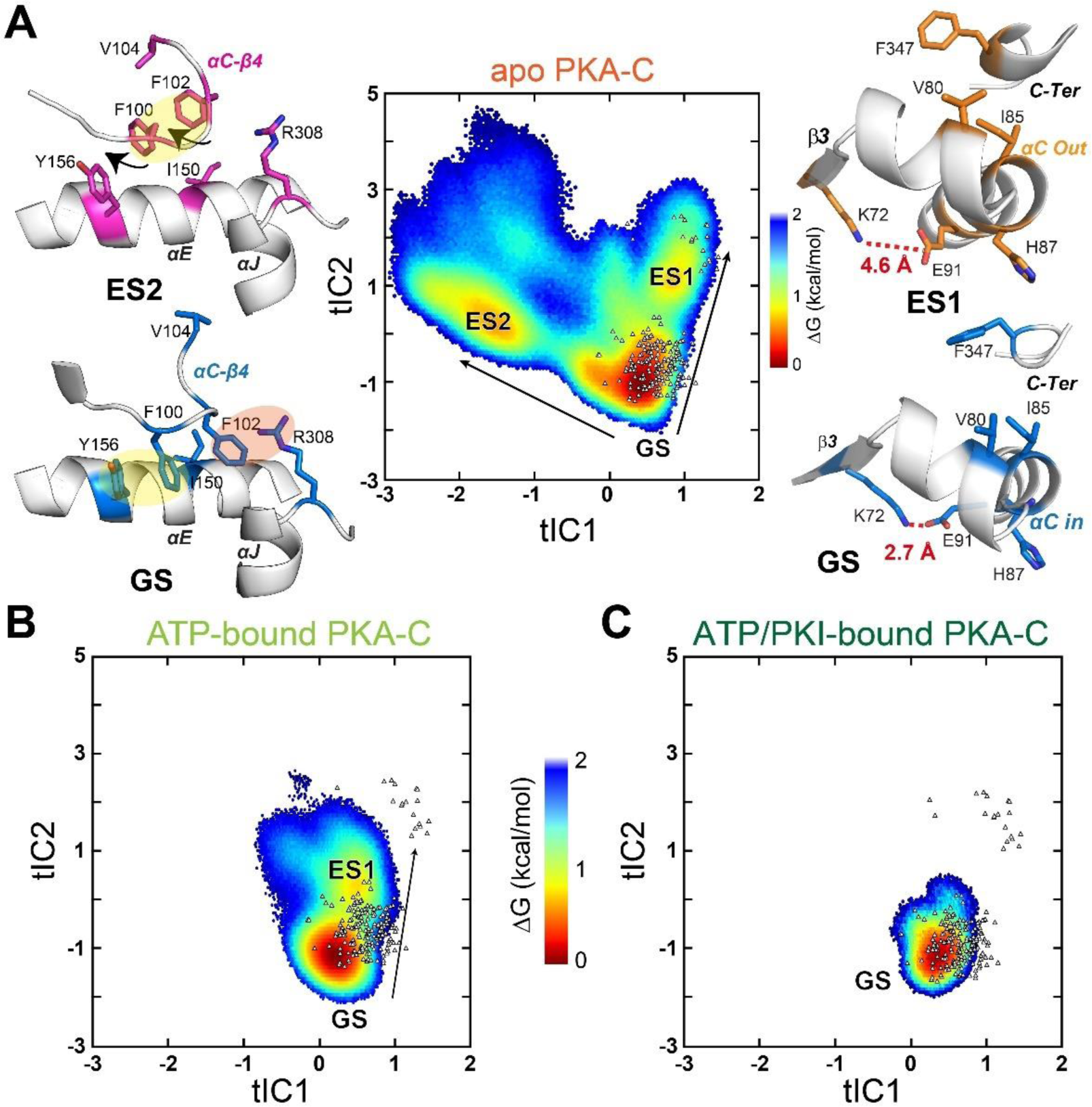
Free energy surfaces and dynamic transitions determined by a Markov State Model (MSM) for apo, ATP-bound, and ATP/PKI-bound PKA-C. (**A**) Free energy landscape projected along the first two time-lagged independent components (tICs) of apo PKA-C, featuring three basins, GS, ES1, and ES2. The transition from GS to ES1 (arrow) highlights the changes around the αB-αC loop, with the disruption of the K72-E91 salt bridge and the PIF pocket (V80-I85-F347) hydrophobic interactions. The GS to ES2 transition (arrow) displays the rearrangement of the hydrophobic packing around the αC-β4 loop. (**B**, **C**) Free energy surfaces projected along the first two tICs for the ATP-bound and ATP/PKI-bound PKA-C, respectively. Known crystal structures for the three forms are indicated by small white triangles.

Moreover, in the apo form, the αC-β4 loop is quite dynamic due to transient hydrophobic interactions involving F100 and V104, as well as W222 and the APE motif (A206 and P207) (**Figure 4A**). The binding of both nucleotide and PKI rigidifies the residues near F100 and V104, *e.g.*, V103 and F185 (**Figure 4A**) and makes more persistent several electrostatic interactions essential for catalysis (D166-N171, K168-T201, and Y204-E230), which are transient in the apo PKA-C. Finally, we used kinetic Monte Carlo sampling to characterize the slow transition between different states.^42^ In the GS state of the apo PKA-C, F100 and F102 adopt *trans* configurations that stabilize the interactions with the αE and αJ helices, and, together with the nucleotide, they lock the αC-β4 loop in an active state (**Figure 3A**). The GS to ES1 transition features the disruption of the K72-E91 salt bridge, typical of the inactive kinase. In contrast, the GS to ES2 transition involves a 120° flip of the F100 aromatic group that interacts with V104, a conformation found only in the catalytically uncommitted apoenzyme (**Figure 3A**). Interestingly, the GS to ES2 transition involves a concerted disruption of the D166-N171 and K168-T201 electrostatic interactions and the destabilization of the packing between W222 and the APE motif (A206 and P207) required for substrate recognition (**Figure 3C**). Overall, these calculations show that conformational transitions from the active GS to ES1 and ES2 represent two independent pathways toward inactive states of the kinase.

**Figure 4.**
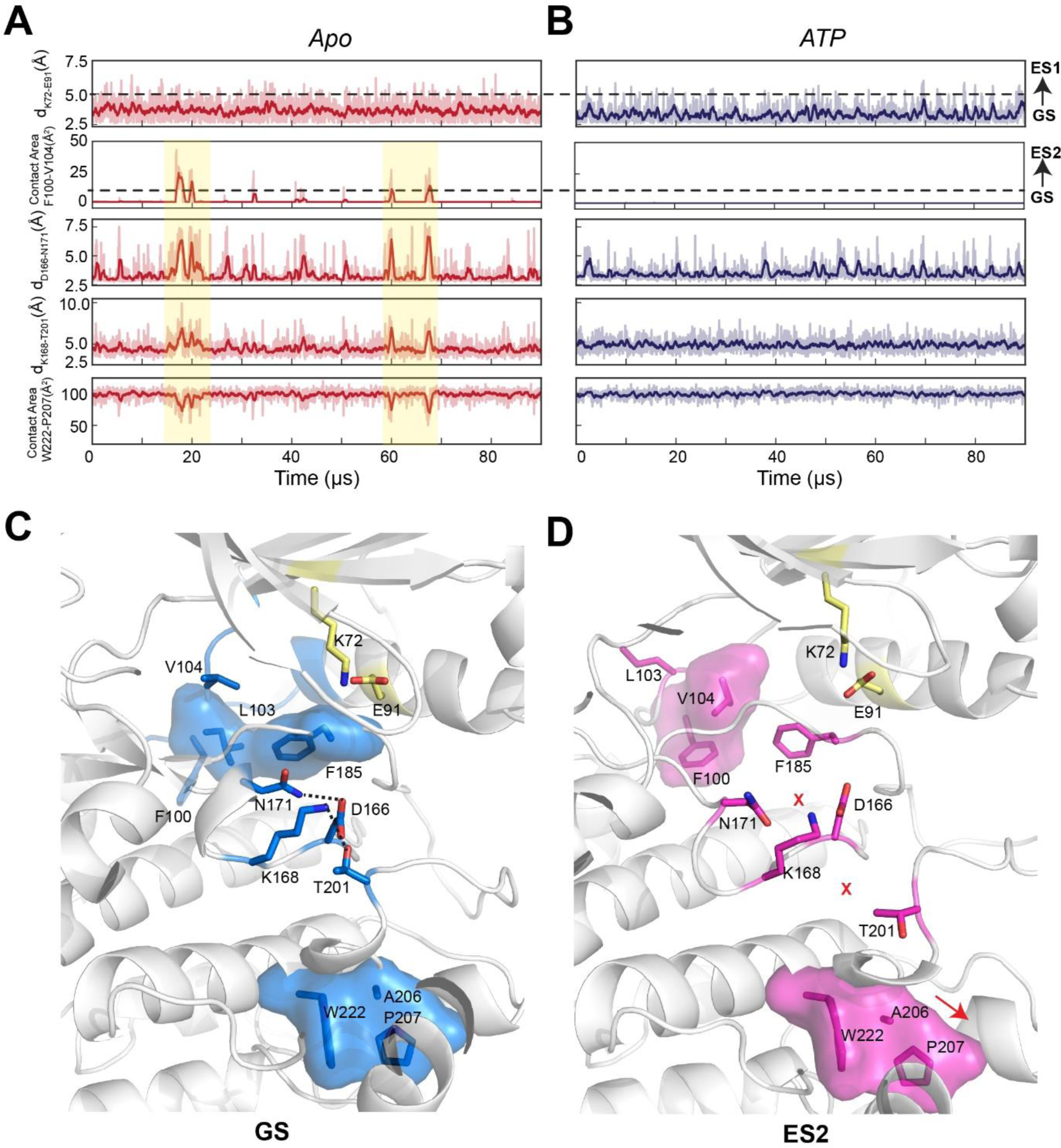
Conformational transitions of apo and ATP-bound PKA-C from GS to ES1 and ES2 states as shown by the kinetic Monte Carlo trajectories. (**A**, **B**) Time course of the structural transitions from GS to ES1 GS to ES2 for apo and ATP-bound PKA-C, respectively. The GS to ES1 transition is characterized by the disruption of the K72-E91 salt bridge and occurs frequently for both PKA-C forms. In contrast, the structural transition from GS to ES2 occurs only for the apo PKA-C, and it features the interactions between F100 and V104 that cause allosteric changes between D166-N171, K168-T201, and W222-A206-P207. The dark color traces indicate the moving averages calculated every 10 frames. (**C**) Structural snapshot of the GS conformation showing that the key catalytic motifs are poised for phosphoryl transfer. (**D**) Structural snapshot of the ES2 conformation with a disrupted configuration of key catalytic motifs typical of inactive kinase.

### Direct correspondence between the conformationally excited states identified by MD simulations and NMR data

NMR CPMG relaxation dispersion and CEST experiments performed on the apo PKA-C revealed the presence of conformationally excited states for several residues embedded into the hydrophobic core of the enzyme.^27^ The chemical shifts of one of these states follow the previously observed open-to-closed transitions of the enzyme,^41^ whereas another excited state observable in the CEST experiments does not follow the same trend. We surmised that this new state may represent an alternate inactive state.^27^ Since ES2 features a disrupted hydrophobic packing of the αC-β4 loop and several of the methyl groups displaying the excited state in the CEST experiments are near the αC-β4 loop, we compared the CS differences of the methyl group from CEST experiments^27^ (Δω^Exp^) with those from the MD simulations (Δω^Pred^). We analyzed the methyl groups of L103, V104, I150, L172, and I180, sampling more than 500 snapshots as representative conformations of the kinase in the ES2 and GS states (**Figure 3 – figure supplement 3**). We then computed the distribution of the methyl ^13^C CS using ShiftX2.^43^ For Val104 and Ile150, these calculations yielded Δω^Pred^ values of 0.87 ± 0.02 and 0.85 ± 0.02 ppm between the ES2 and GS conformations, respectively, which are in good agreement with Δω^Exp^ (1.10 ± 0.06 and 0.94 ± 0.09 ppm) obtained from the CEST experiments or fitting the CPMG dispersion curves (**Figure 5A-B**). The Δω^Pred^ values and the directions of the CS changes obtained for the remainder three sites are also in good agreement with the CEST profiles (**Figure 5 – figure supplement 1**). **Figure 4C** shows a linear fitting of Δω^Pred^ and Δω^Exp^ with a slope of 0.86 and R^2^ of 0.82. The latter supports the hypothesis that the excited state observed by NMR may correspond to the simulated structural ensemble of the ES2 basin. Finally, MSM estimates a population of ES2 of 6 ± 2%, which is consistent with the 5 ± 1% population found by fitting the NMR data.^40^ Altogether, the combination of NMR and MD simulations support the presence of an alternate inactive state that features disrupted hydrophobic interactions near the αC-β4 loop that perturb the anchoring with the αE helix, and in turn, the structural couplings between the two lobes of the kinase.

**Figure 5.**
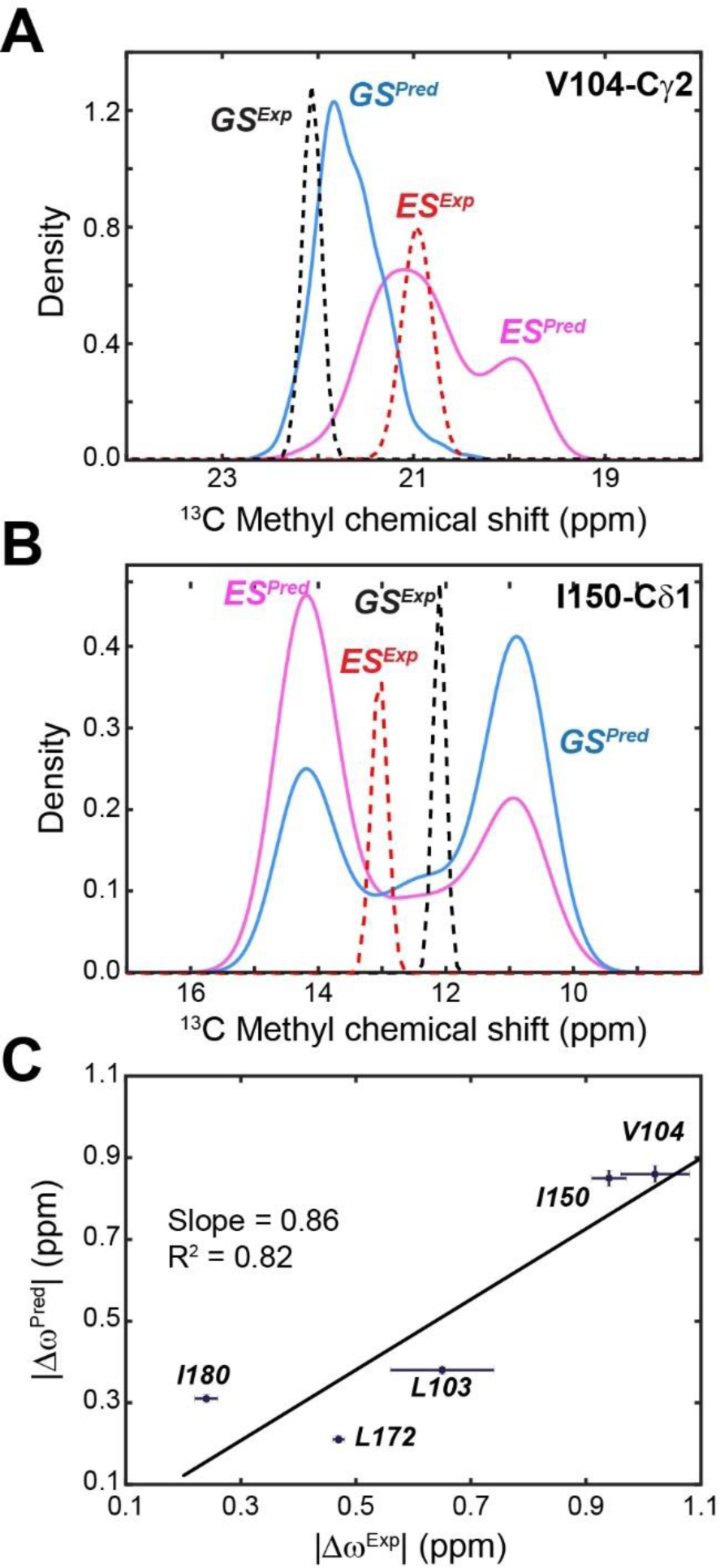
Comparison of experimental versus calculated ^13^C chemical shifts of methyl groups of PKA-C. (**A**) Experimental and calculated CS for Val104-Cγ1 of apo PKA-C. The GS is in blue and the ES in magenta. (**B**) Corresponding CS profiles for Ile150-Cδ1. The experimental CS is shown in dotted lines for GS (black) and ES (red). (**C**) Correlation between predicted | Δω^Pred^| and experimental | Δω^Exp^| CS differences for methyl groups near the αC-β4 loop. The fitted linear correlation has a slope of 0.86 and R^2^ of 0.82.

### The F100A mutation disrupts the allosteric network of the kinase

The above analysis suggests that the destabilization of the αC-β4 loop may promote the GS to ES2 transition of the kinase. To validate this hypothesis, we generated the F100A mutant seeking to increase the flexibility of the αC-β4 loop. Starting from the coordinates of the X-ray structure of the ternary complex (PDB ID: 4WB5), we simulated the F100A dynamics in an explicit water environment. First, we performed a short equilibration using classical MD simulations. During 1 μs of MD simulations, the αC-β4 loop of the F100A mutant undergoes a significant motion as manifested by increased values of the backbone root mean square deviation (rmsd) and a conformational change (flipping) of the F102 side chain (**Figure 6A**). This region in PKA-C^WT^ adopts a stable β-turn, with a persistent hydrogen bond between the backbone oxygen of F100 and the amide hydrogen of L103. In contrast, the hydrogen bond in F100A is formed more frequently between A100 and F102, resulting in an average γ-turn conformation (**Figure 6B**). Such local rearrangement not only disrupts the hydrogen bond between N99 and Y156, altering the interactions between the αC-β4 loop and the αE helix but also destabilizes the cation-π interaction between F102 and R308 at the C-tail (**Figure 6B – supplementary figure 1**)

**Figure 6.**
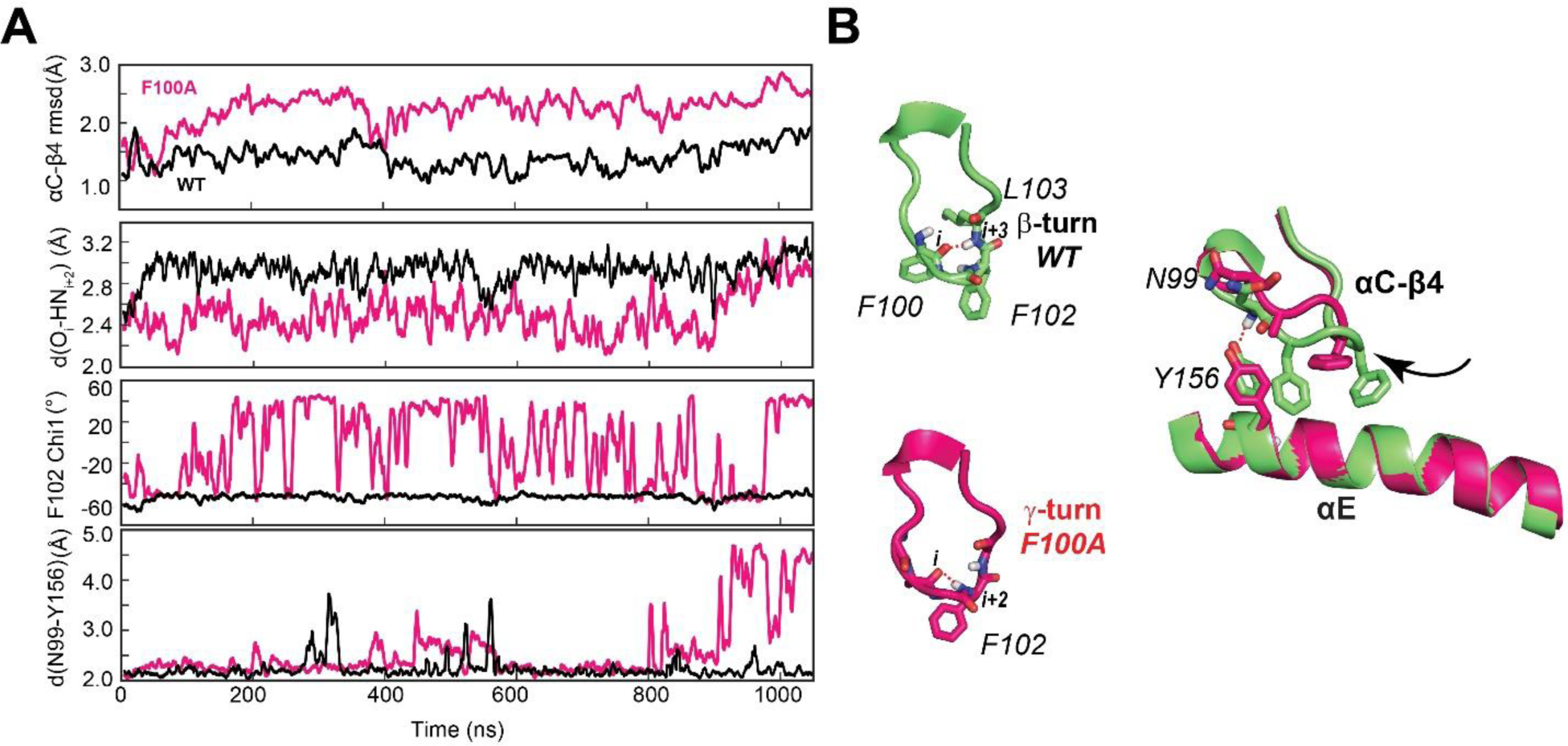
F100A mutation increases the dynamics of the αC-β4 loop, perturbing the local hydrophobic packing and its anchoring to the αE helix. (**A**) Time series of the αC-β4 loop dynamics, H-bond occurrence for the β− and γ-turns, F102 χ_1_ angle, and N99 and Y156 for WT (black) and F100A (red) in the ATP-bound state. (**B**) Representative structural snapshots showing the β-turn conformation for PKA-C^WT^ (green) and γ-turn for PKA-C^F100A^ (magenta).

The structural changes of the αC-β4 loop caused by the F100A mutation propagate through the protein core across the R spine, C spines, and shell residues (**Figure 7A**), and alter the response of the kinase to nucleotide binding. In the PKA-C^WT^, ATP binding shifts the conformational ensemble of the kinase toward an intermediate state competent to substrate binding, as represented by the changes in the population densities of the hydrophobic core residues as a function of the rmsd (**Figure 7B**). For F100A, the population of the C spine follows the same trend of the WT kinase, with the shell residues already populating the intermediate state. In contrast, the R spine residues fail to adopt a competent state (**Figure 7B**). The latter is due to the perturbation of hydrophobic packing of L95 and L106 in the R spine, and V104 in the shell near the mutation site. Using the lowest principal components, we also analyzed the global dynamic response of the kinase to ATP binding. Not only does F100A change the breathing mode of the two lobes (PC1), but it also alters the shearing motion (PC2) of the binary complex, emphasizing the importance of this allosteric site for the inter-lobe communication (**Figure 7C-E**).

**Figure 7.**
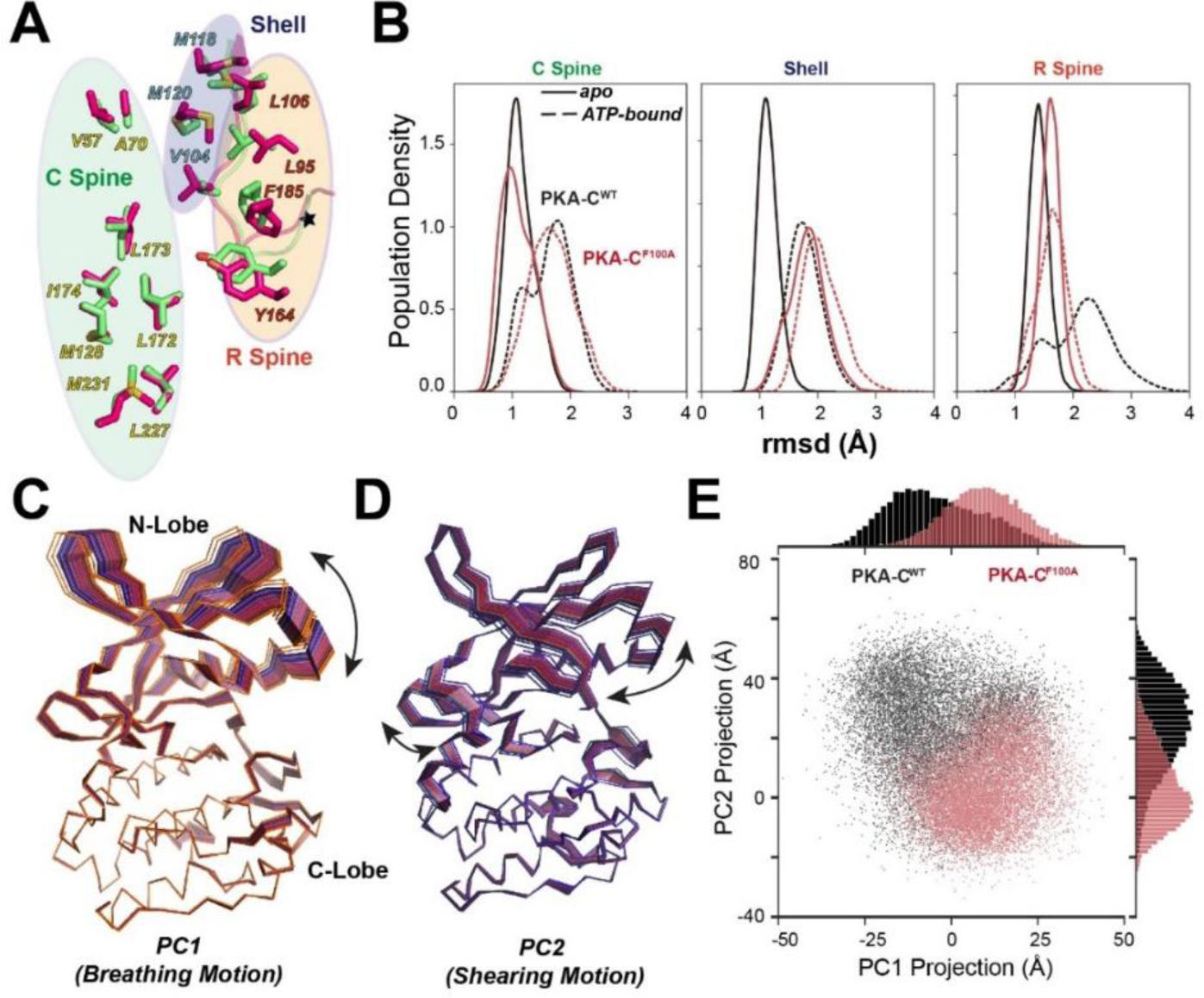
Structural responses to ATP binding of PKA-C^F100A^ mutant. (**A**) Superposition of the hydrophobic cores (C spine, R spine, and shell residues) for PKA-C^WT^ (lime) and PKA-C^F100A^ (hot pink), highlighting the structural perturbations of the R spine and shell residues. (**B**) Structural perturbation upon ATP binding for the hydrophobic core residues of PKA-C^WT^ and PKA-C^F100A^ shown as changes in the population densities vs. rmsd (**C, D**) First (PC1) and second (PC2) principal components describing breathing and shearing motions of the two lobes. (**E**) 2D projections and distributions of PC1 and PC2 for PKA-C^WT^ and PKA-C^F100A^.

To analyze the internal correlated motions of the kinase, we further analyzed the MD trajectories of both wild-type and mutant enzyme using mutual information (**Figure 8**).^44^ This method identifies correlated changes of backbone and side chain rotamers in proteins and detects clusters of residues that are responsible for intramolecular communication between active sites.^44^ For PKA-C^WT^, we observed numerous strong correlations that are clustered within each lobe and across the enzyme, connecting key motifs, such as the Gly-rich loop, αC-β4 loop, activation loop, and C-terminal tail (**Figure 8A**). These correlation patterns are reminiscent of the chemical shift correlations experimentally found for PKA-C^WT^,^20^ which constitute the hallmark of the dynamically-committed state.^35,36^ In contrast, the mutual information matrix for the F100A mutant displays weaker and interspersed correlations across the enzyme. New correlations also are present across the helical C-lobe suggesting a rewiring of the internal allosteric communication (**Figure 8B**). A possible explanation is that the increased motion caused by the elimination of the F100 aromatic side chain at the αC-β4 loop propagates throughout the entire hydrophobic core. This structural reorganization causes the F100A kinase to adopt a dynamically uncommitted state.^35^ Taken together, the simulations of F100A suggest that perturbations of the aC-b4 loop increase the local flexibility and break the structural connection between the two lobes, disrupting the correlated breathing/shearing motions highlighted in previous MD simulation studies^45,46^. Although the single F100A mutation does not drive the kinase into a completely inactive state (ES2), it is sufficient to abolish the structural couplings between the two lobes.

**Figure 8.**
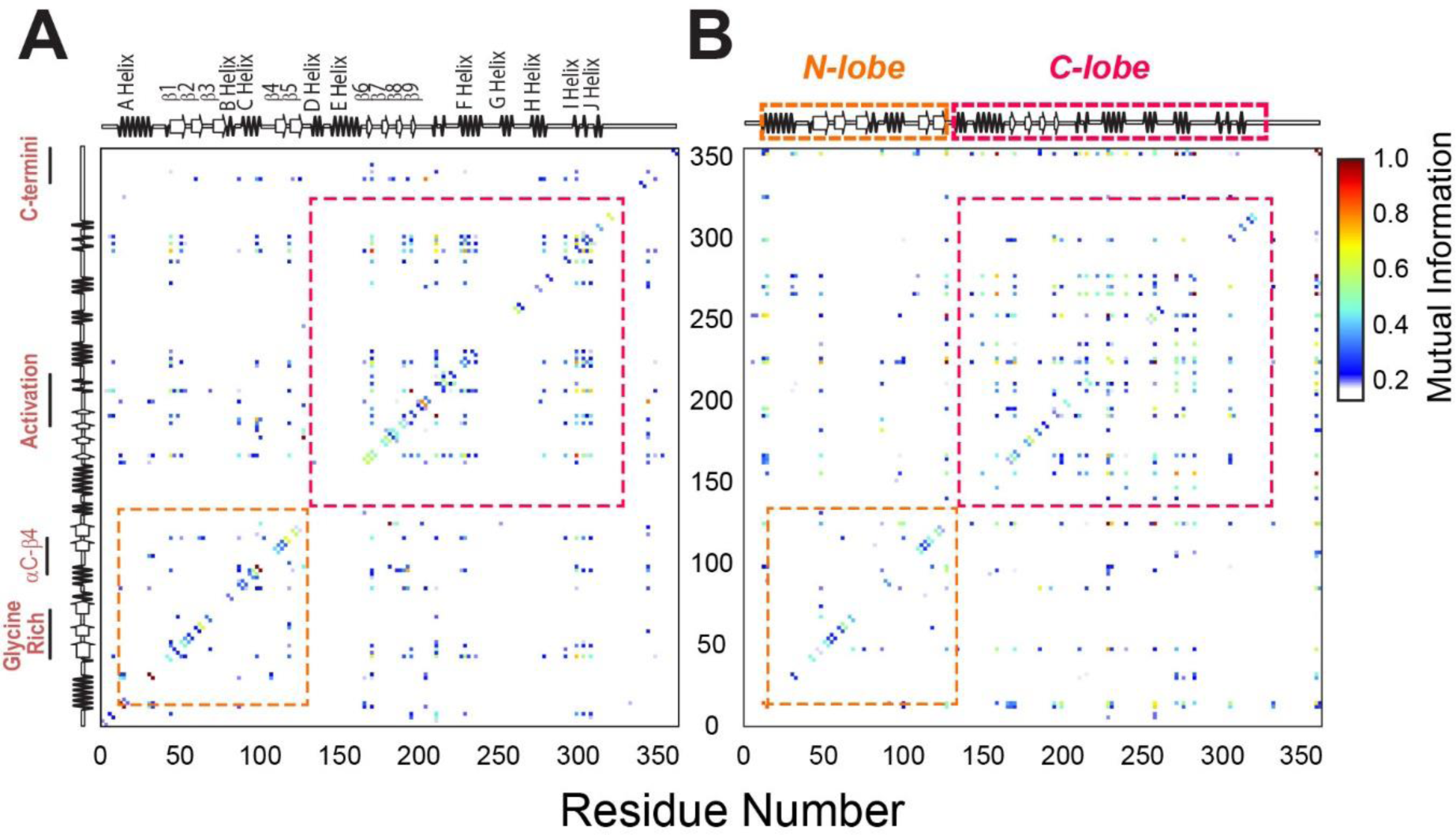
Mutual information analysis of backbone and side chain rotamers of ATP-bound PKA-C^WT^ and PKA-C^F100A^. (**A**) Mutual information matrix for PKA-C^WT^ showing well-organized clusters of interactions within each lobe and distinct inter-lobe communication typical of an active kinase. (**B**) Mutual information matrix for PKA-C^F100A^ revealing an overall reorganization of the allosteric network caused by the disruption of the hydrophobic core.

### Effects of the F100A mutation on the catalytic efficiency and binding thermodynamics of PKA-C

Based on the MD simulations of the F100A mutant, we hypothesized that the disruption of the structural coupling between the N-lobe and C-lobe would affect the binding cooperativity between nucleotide (ATPγN) and pseudosubstrate inhibitor (PKI_5-24_). To test this hypothesis, we expressed, purified, and evaluated the catalytic efficiency for the WT and F100A kinases by carrying out steady-state coupled enzyme assays^47^ using the standard substrate Kemptide.^48^ F100A showed only a slight increase in *K_M_* and *V_max_* compared to PKA-C^WT^, resulting in a small reduction of the catalytic efficiency (*k_cat_/K_M_* = 0.50 ± 0.04 and 0.41 ± 0.08 for PKA-C^WT^ and PKA-C^F100A^, respectively; **Supplementary table 2**). We then performed ITC^49^ to obtain Δ*G*, Δ*H*, *-T*Δ*S*, *and K_d_*, and determine the cooperativity coefficient (σ) for ATPγN and PKI_5-24_ binding. We first analyzed the binding of ATPγN to the apo PKA-C^F100A^ and, subsequently, the binding of PKI_5-24_ to the ATPγN-saturated PKA-C^F100A^ (**Supplementary Table 3**). We found that PKA-C^WT^ and PKAC^F100A^ have similar binding affinities for ATPγN (*K_d_* = 83 ± 8 μM and 73 ± 2 μM, respectively). However, in the apo form, F100A showed a 3-fold higher binding affinity for the pseudosubstrate relative to PKA-C^WT^ (*K_d_* = 5 ± 1 μM and 17 ± 2 μM, respectively - **Supplementary Table 3**). Upon saturation with ATPγN, PKA-C^F100A^ displayed a 12-fold reduction in binding affinity for PKI_5-24_, resulting in a σ of ∼3, a value significantly lower than the WT enzyme (σ greater than 100). These data support the predictions of MD simulations and mutual information that ATP binding to F100A does not promote a conformational state fully competent with substrate binding.

### NMR mapping of nucleotide/pseudosubstrate binding response

To elucidate the atomic details of the disrupted structural coupling and binding cooperativity for PKA-C^F100A^, we used solution NMR spectroscopy mapping its response to the nucleotide (ATPγN) and pseudosubstrate (PKI_5-24_) binding. Specifically, we monitored the ^1^H and ^15^N chemical shift perturbations (CSPs, Δδ) of the amide fingerprints for the binary (PKA-C^F100A^/ATPγN) and ternary (PKA-C^F100A^/ATPγN/PKI_5-24_) complexes and compared them with the corresponding complexes of PKA-C^WT^ (**Figure 9** and **Figure 9 – figure supplement 1**). The interaction with ATPγN causes a dramatic broadening of several amide resonances throughout the structure of PKA-C^F100A^, suggesting the presence of an intermediate conformational exchange as previously observed.^50^ For the remainder of the amide peaks, both PKA-C^WT^ and PKA-C^F100A^ exhibit similar CSP profiles upon binding ATPγN (**Figure 9A-B**), although the extent of the CS changes is significantly attenuated for F100A (**Figure 9 – figure supplement 1B**). This reduction is apparent for residues in the N-lobe (β2-β3 region), around the mutation site (β4), and at the C-terminal tail. A similar pattern emerges upon binding PKI_5-24_ to the ATPγN-saturated PKA-C^F100A^, where a substantial decrease in CSP is observed for residues of the C-lobe localized near the motifs critical for substrate-binding (*i.e.*, αE, αF, and αG helices, **Figure 9C-D**).

**Figure 9.**
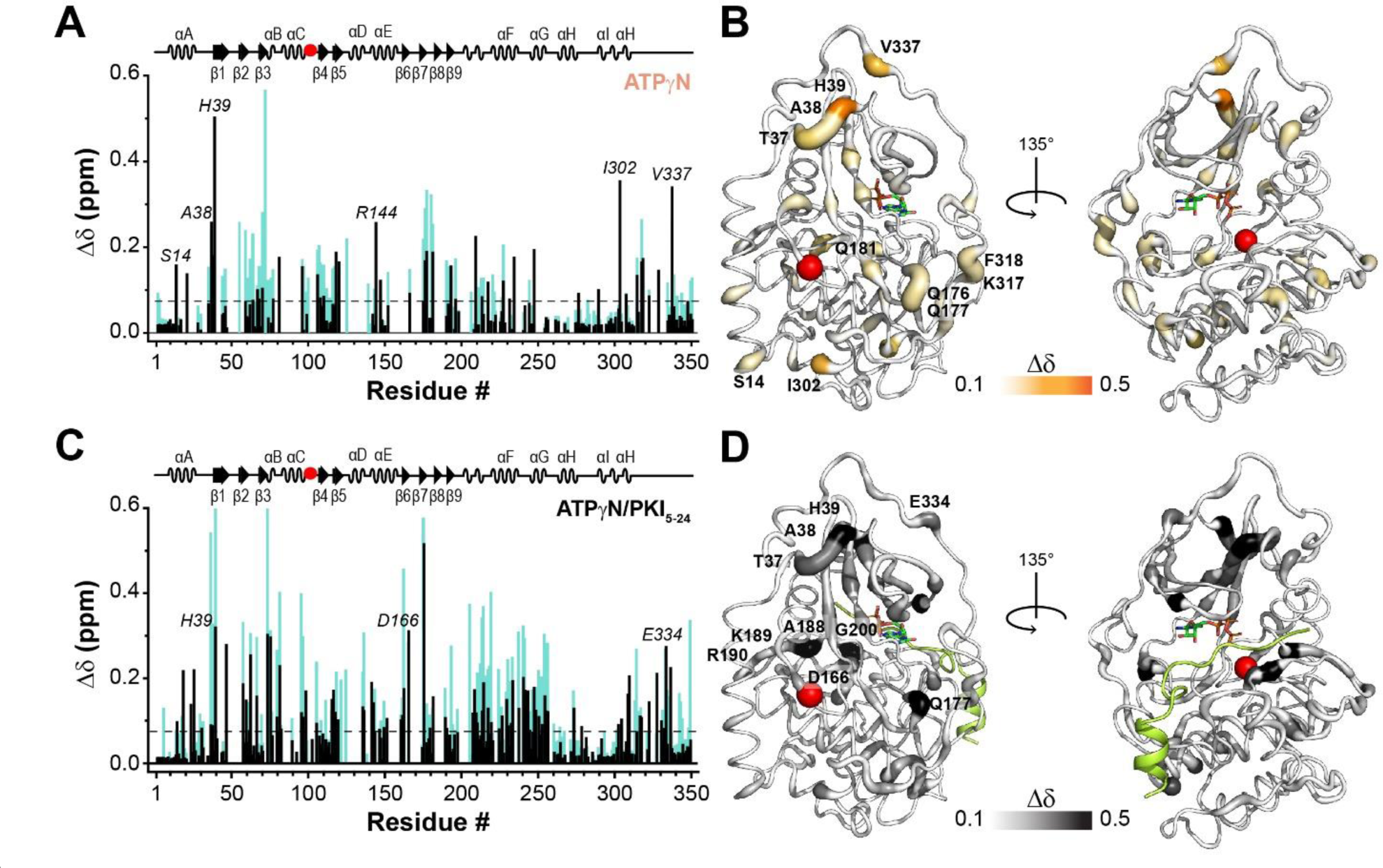
NMR map of the structural response of PKA-C^F100A^ to nucleotide and PKI binding. (**A**) Comparison of the chemical shift perturbation (CSP) of the amide resonances for PKA-C^F100A^ (black) and PKA-C^WT^ (cyan) upon ATPγN binding. The dashed line indicates one standard deviation from the average CSP. (**B**) CSPs of PKA-C^F100A^/ATPγN amide resonances mapped onto the crystal structure (PDB: 4WB5). (**C**) Comparison of the CSPs of the amide resonances for PKA-C^F100A^ and PKA-C^WT^ upon binding ATPγN and PKI_5-24_ (black). (**D**) CSPs for the F100A/ATPγN/PKI complex mapped onto the crystal structure (PDB: 4WB5). To define the allosteric network of the kinase upon binding nucleotides and substrate, we examined the CS using CHEmical Shift Covariance Analysis (CHESCA),^22,23,27^ a statistical method that identifies correlated responses of residue pairs to a specific perturbation (*i.e.*, ligand binding, mutations, *etc.*). CHESCA works under the assumption that pairwise correlated CS changes of residues identify possible intramolecular allosteric networks.^52,53^ For PKA-C, we found that coordinated structural rearrangements, as identified by CHESCA, are directly related to the extent of binding cooperativity.^22,23,27^ Therefore, we compared the CHESCA maps for PKA-C^WT^ and PKAC^F100A^ for four different states: *apo*, ATPγN-, ADP-, and ATPγN/PKI_5–24_-bound. For PKA-C^F100A^, the CHESCA matrix exhibits sparser and more attenuated correlations (*i.e.*, lower correlation co-efficient value) relative to PKA-C^WT^ (**Figure 10A**). Although many inter-lobe correlations are still present for F100A, several other correlations in specific structural domains such as the αG-, αH-, and αI-helices are absent or attenuated. For instance, the F100A mutation does not display cor-relations between the αA-helix and the C-terminal tail that constitute a critical “complement to the kinase core”.^54^ We also utilized CHESCA to assess the allosteric communication among the PKA-C communities as defined by McClendon *et al*.^55^ The CHESCA community map for PKA-C^WT^ shows strong correlations across the enzyme, especially for structurally adjacent communities and at the interface between the two lobes (see for instance the correlations among *ComA*, *ComB*, *ComC*, *ComE*, and *ComH*) (**Figure 10B-C**). For F100A, the CHESCA community map shows that the cross-talk between the nucleotide-binding (*ComA*) and positioning of αC-helix (*ComB*) communities, as well as the R-spine assembly (*ComC*) and the activation loop (*ComF*) communities are preserved (**Figure 10B-C**). However, the correlations between *ComE*, responsible for stabilizing the C spine, and *ComC*, involved in the assembly of the R spine, are absent. Similarly, the long-range correlations between the C and R spines (*i.e.*, *ComD* with *ComC*) are missing. Finally, several correlations between *ComF1*, *ComG*, and *ComH* are no longer present. These communities orchestrate substrate recognition and R subunits binding. Overall, the CHESCA analysis for PKA-C^F100A^ suggests that the reduced degree of cooperativity we determined thermodynamically corresponds to a decrease in coordinated structural changes upon ligand binding. The latter is apparent from the loss of correlated structural changes among the structural communities, including the hydrophobic spines, substrate binding cleft, and the docking surface for PKA interactions with other binding partners.

**Figure 10.**
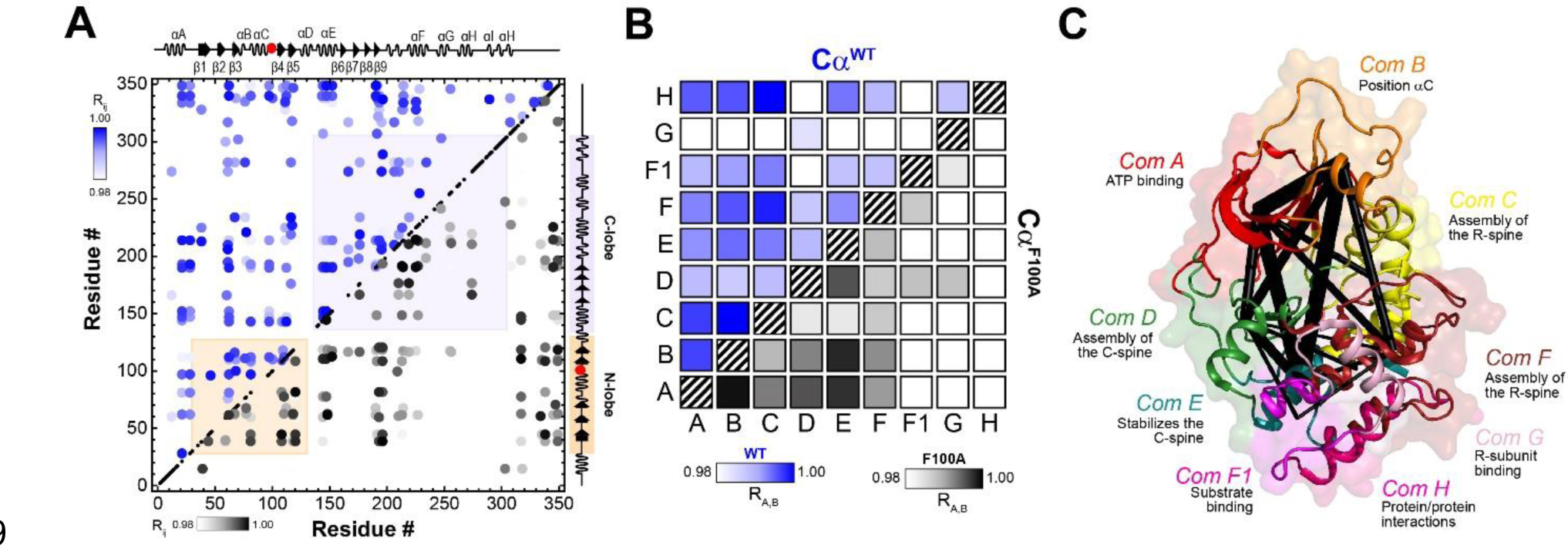
Correlated chemical shift changes reveal the uncoupling of the intramolecular allosteric network in PKA-C^F100A^. (**A**) Comparison of the CHESCA matrices obtained from the analysis of the amide CS of PKA-C^WT^ (blue correlations) and PKA-C_F100A_ (black correlations). The correlations coefficients (R_ij_) were calculated using the apo, ADP-bound, ATPγN-bound, and ATPγN/PKI_5-24_-bound states. For clarity, only correlation with R_ij_ > 0.98 are displayed. For the enlarged CHESCA map of F100A see **Figure 10 – figure supplement 1.** The data for the PKA-C^WT^ matrix were taken from Walker *et al.*^22^. (**B**) Community CHESCA analysis of PKA-C^WT^ (blue correlations) and PKA-C^F100A^ (black correlations). Only correlations with R_A,B_ > 0.98 are shown. Spider plot showing the extent of intramolecular correlations identified by the community CHESCA analysis for PKA-C^F100A^ mapped onto the crystal structure (PDB: 4WB5). The thickness of each line in the spider plot indicates the extent of coupling between the communities.

To determine the global response to ligand binding for WT and F100A, we analyzed the chemical shifts of the amide fingerprints of the two proteins using the COordiNated ChemIcal Shifts bEhavior (CONCISE).^51^ CONCISE describes the overall changes of the protein fingerprint resonances by providing the probability density (population) of each state along the conformational equilibrium for binding phenomena that follow linear CS trajectories.^51^ For PKA-C^WT^, both nucleotide and pseudosubstrate binding shift the overall population of the amides from an open to an intermediate and a fully closed state (**Figure 9 – figure supplement 2**). A similar trend is observed for PKA-C^F100A^, though the probability densities are broader, indicating that the amide resonances follow a less coordinated response.^51^ Also, the maximum of the probability density for the ternary complex is shifted toward the left, indicating that the mutant adopts a more open structure than the corresponding wild-type enzyme. Overall, the shapes and the positions of the probability distributions of the resonances suggests that several residues do not respond in a coordinated manner to the nucleotide binding, and PKI_5-24_ shifts the conformation of the kinase toward a partially closed state, which explains the loss in binding cooperativity as previously observed.^21-^

## DISCUSSION

Our structural and dynamic studies suggest that the binding cooperativity of PKA-C originates from the allosteric coupling between the nucleotide-binding pocket and the interfacial region between the two lobes, which harbors the substrate binding cleft.^21–23^ The latter has been emphasized for other kinases such as Src and ERK2.^56,57^ The biological significance of this intramolecular communication has emerged from our recent studies on pathological mutations situated in the activation loop of PKA-C and linked to Cushing’s syndrome.^21–23^ These mutations drastically reduce substrate binding affinity, and more importantly, disrupt the communication between the two ligand binding pockets of the kinase.^21–23^ Interestingly, the E31V mutation distal from the active site and also related to the Cushing’s syndrome, displays disfunction similar to the other orthosteric mutations, suggesting that it is possible to modulate substrate recognition allosterically.^23^ Indeed, these mutations are connected to allosteric nodes that, once perturbed, radiate their effects towards the periphery of the enzyme interrupting the coordinated dynamic coupling between the two lobes.^23^ Interestingly, these mutations do not prevent Kemptide phosphorylation; rather, they cause a loss of substrate fidelity with consequent aberrant phosphorylation of downstream substrates.^10,58–62^ Additionally, recent thermodynamic and NMR analyses of PKA-C binding to different nucleotides and inhibitors demonstrated that it is possible to control substrate binding affinity by changing the chemistry of the ligand at the ATP binding pocket.^22^ Altogether, these studies put forward the idea that substrate phosphorylation and cooperativity between ATP and substrates may be controlled independently.

By integrating our NMR data with RAM simulations and MSM, we comprehensively mapped the free energy landscape of PKA-C in various forms. We found that the active kinase unleashed from the regulatory subunits occupies a broad energy basin (GS) that corresponds to the conformation of the ternary structure of PKA-C with ATP and pseudosubstrate (PKI_5-24_), exemplifying a catalytic competent state poised for phosphoryl transfer.^15^ We also identified two orthogonal conformationally excited states, ES1 and ES2. While ES1 corresponds to canonical inactive kinase conformations, the ES2 state was never observed in the crystallized structures. Our previous CEST NMR measurements suggested the presence of an additional sparsely populated state that, at that time, we were unable to characterize structurally.^27^ The chemical shifts of this state did not follow the opening and closing of the kinase active cleft and were assigned to a possible alternate inactivation pathway.^27^ These new simulations and MSM show that this sparsely populated state may be attributed to the transition from GS to ES2, which features a disruption of hydrophobic packing, with a conformational rearrangement for the αC-β4 loop that causes a partial disruption of the hydrophobic R spine. These structural changes rewire the allosteric coupling between the two lobes, as shown by mutual information analysis. A single mutation (F100A), suggested by our simulations, promotes the flip of the αC-β4 loop and reproduces the hypothesized structural uncoupling between the two lobes of PKA-C. We experimentally tested the effects of the F100A mutation and found that it prevents the enzyme from adopting a conformation competent for substrate binding, resulting in a drastic reduction of the cooperativity between ATP and substrate. These NMR data further support our working model, showing that if the inter-lobe communication is interrupted, the binding response of the kinase is attenuated. Indeed, our investigation further emphasizes the integral role of the kinase hydrophobic core and demonstrates that alterations in the spines and shell residues may lead to a dysfunctional kinase by either preventing phosphorylation (see V104G and I150A mutations)^40^ or by disrupting the binding cooperativity as for F100A.

The αC-β4 loop is a regulatory element present in all EPKs and its importance has been stressed in bioinformatics studies^63^ and supported by computational work.^55^ F100 and F102 in the αC-β4 loop are at intersections of various structural communities and constitute a critical hydrophobic node that anchors the N- to the C-lobe. Also, studies on EGFR and ErbB2 kinases led to the hypothesis that the αC-β4 loop may act as a molecular brake ^64,65^ or an autoinhibitory switch.^66^ Therefore, it is not surprising that activating mutations and in-frame insertions in the αC-β4 loop are frequently found in kinase-related cancers and somatic oncogenic mutations such as P101S, P101L, and L103F.^67^ Also, an elegant study by Kannan and coworkers emphasized the role of the αC-β4 loop in dimerization and aberrant activation of EGFR by insertion mutations.^28^ Moreover, in a recent paper Zhang and co-workers found a similar behavior for the HER2 exon 20 insertions.^30^ These latter two studies propose that in-frame insertions alter the conformational landscape of the kinase rigidifying the structure around the αC-β4 loop and restricting the kinase conformational ensemble in a constitutively active state. Remarkably, in the case of EGRF insertion mutations, the activation of the kinase occurs in a length-dependent manner. ^28^ Additionally, these human mutations display a gradual response to drugs, which can be exploited for designing selective inhibitors against EGFR pathological mutants.^68^

The data presented here show that it is possible to abolish the binding cooperativity of a kinase by turning the dial in the opposite direction, *i.e.*, increasing the flexibility of the αC-β4 loop and disconnecting the allosteric network between the N- and C-lobes. The identification of this new, partially inactivating pathway provides further understanding of how to control the dynamics and function of kinases.

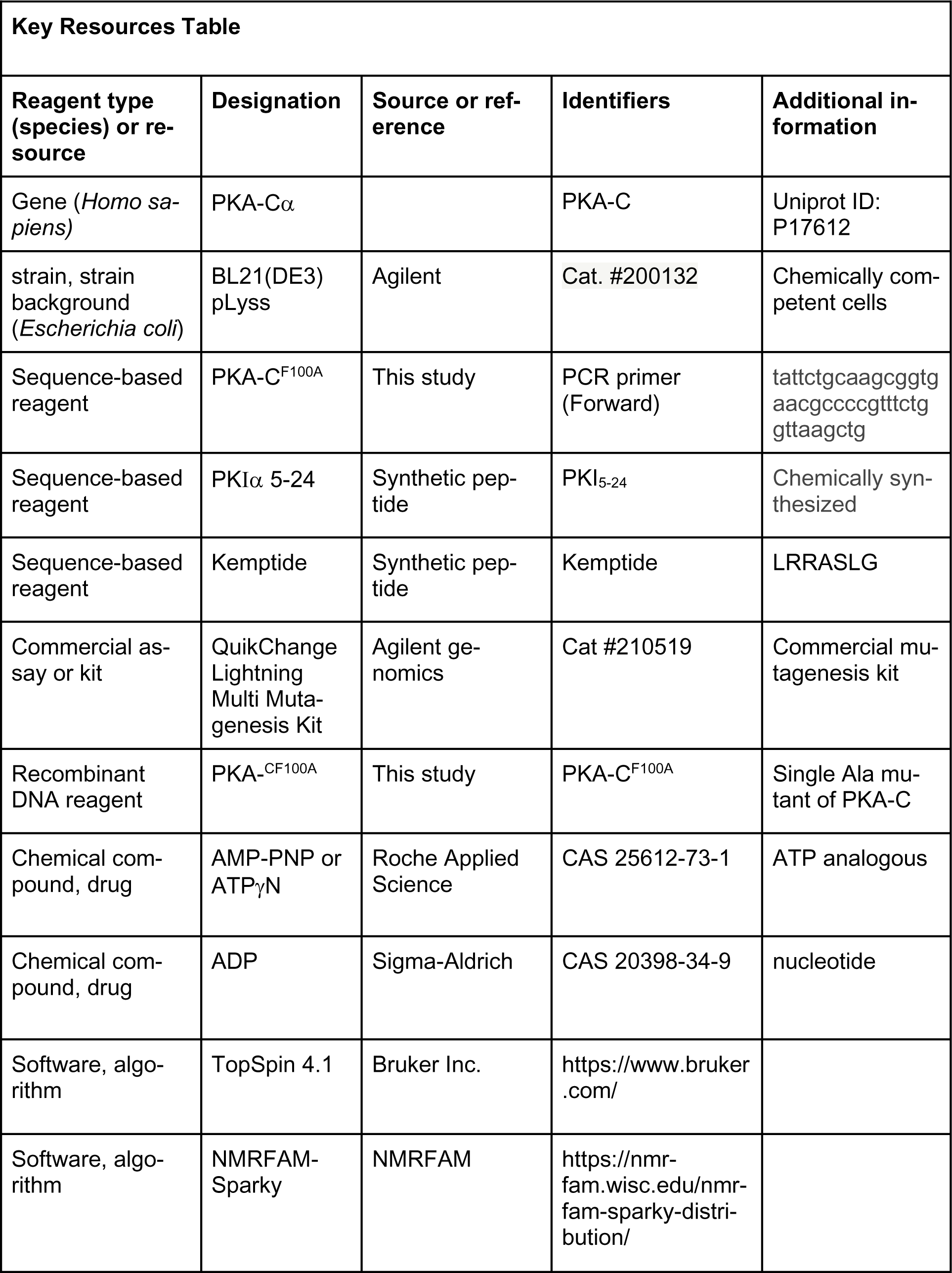
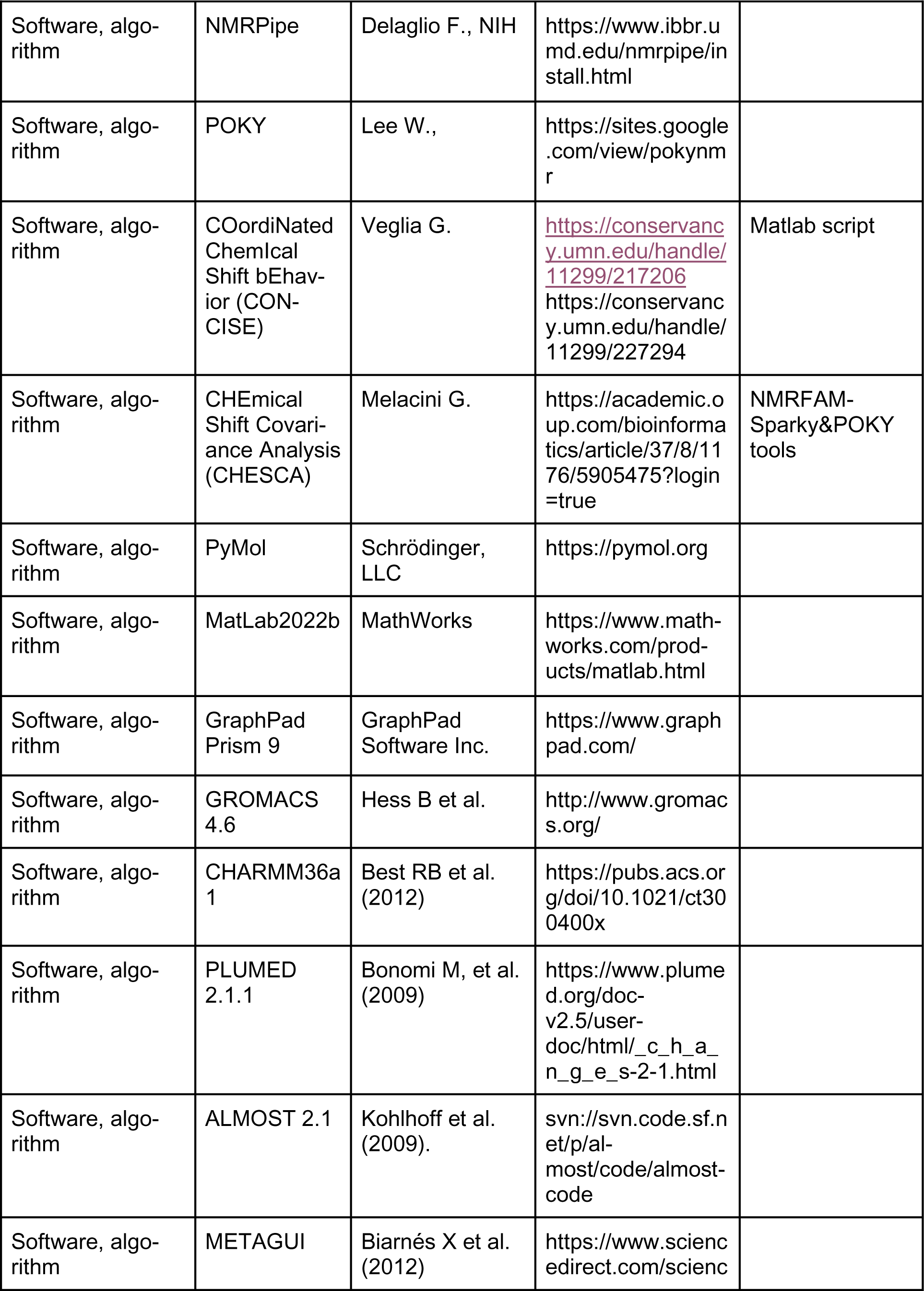
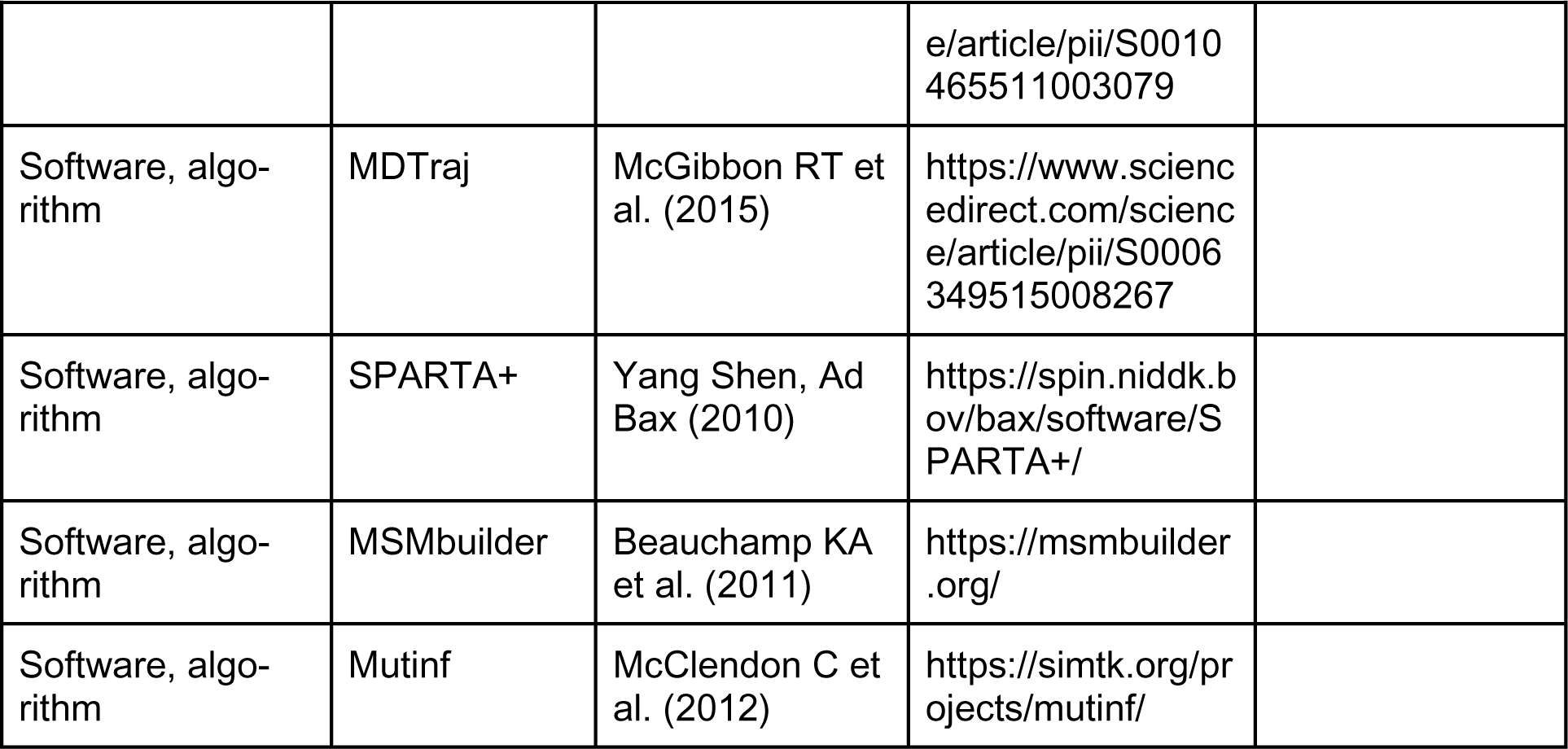

## MATERIAL AND METHODS

### Replica-averaged metadynamics (RAM) simulations

#### I. System setup

As a starting structure, we used the coordinates of the crystal structure of PKA-C^WT^ (PDB ID: 1ATP) and added the missing residues 1-14 at the N terminus. The protonation state of histidine residues was set as previously reported.^69^ The kinase was solvated in a rhombic dodecahedron solvent box using the three-point charge TIP3P model^70^ for water, which extended approximately 10 Å away from the surface of the proteins. Counter ions (K^+^ and Cl^-^) were added to ensure electrostatic neutrality to a final ionic concentration of ∼ 150 mM. All protein covalent bonds were constrained using the LINCS algorithm^71^ and long-range electrostatic interactions were simulated using the particle-mesh Ewald method with a real-space cut-off of 10 Å.^72^ The simulations of apo, binary (one Mg^2+^ ion and one ATP), and ternary (two Mg^2+^ ions, one ATP, and PKI_5-24_ ) forms were performed simultaneously using GROMACS 4.6^73^ with CHARMM36a1 force field.^74^ For F100A, the corresponding residues were mutated through the mutagenesis wizard of Pymol.^75^

#### II. Standard MD simulations

Each system was minimized using the steepest descent algorithm to remove the geometric distortions, and then gradually heated to 300 K at a constant volume over 1 ns using harmonic restraints with a force constant of 1000 kJ/(mol *Å^2^) on heavy atoms for both the kinase and nucleotide. The restraints were gradually released over the following 12 ns of simulations at constant pressure (1 atm) and temperature (300 K). The systems were equilibrated for an additional 20 ns without positional restraints. A Parrinello-Rahman barostat^76^ was used to keep the pressure constant, while a V-rescale thermostat ^77^ with a time step of 2 fs was used to keep the temperature constant. Each system was simulated for 1.05 µs, with snapshots recorded every 20 ps.

#### III. Replica exchange (REX) simulations

Following standard MD simulations, parallel replica exchange (REX) simulations were set up on the apo, binary, and ternary forms of PKA-C. Four replicas were used for each REX simulation, and the initial structures were randomly chosen from the µs-scale unbiased simulations. Chemical shifts of PKA-C for N, CA, CO, CB, and HN from NMR experiments were imposed as restraints based on the following penalty function:

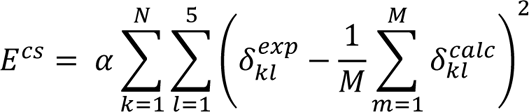

where *α* is the force constant, *k* runs over all residues of the protein, *l* denotes the different backbone atoms, and *m* runs over the four replicas. 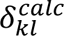 was computed using CamShift, a module of ALMOST-2.1.^78^ The force constant was gradually increased from 0 (unbiased) to 20 (maximum restraints for production) over 50 ns. All other settings were identical to the unbiased simulations. REX simulations were carried out with GROMACS 4.6,^73^ with the replica exchange controlled by the PLUMED 2.1.1 module.^79^ Approximately, 100 ns of REX simulations were further carried out for each replica in the three PKA-C forms.

#### IV. Replica-averaged metadynamics (RAM) simulations

RAM simulations were started from the final structures of REX simulations. The CS restraints were imposed in the same manner as for the REX simulations. Four CVs are chosen to increase the conformational plasticity around the catalytic cores (detailed in **Figure S1**): (CVI) the ψ angles of the backbone of all the loops that are not in contact with ATP (Back-far), (CVII) the ϕ angles of the backbone of all the loops that are in contact with ATP (Back-close), (CVIII) the χ_1_ angles of side-chains of all the loops that are in contact with ATP (Side-close), (CVIV) the radius of gyration calculated over the rigid part (*i.e.*, residues 50-300) of the protein (rgss). Gaussian deposition rate was performed with an initial rate of 0.125 kJ/mol/ps, where the σ values were set to 0.5, 0.2, 0.2, and 0.01 nm for the four CVs, respectively. The RAM simulations were also carried out with GROMACS 4.6 in conjunction with PLUMED 2.1 and ALMOST 2.1, and continued for ∼400 ns for each replica with exchange trails every 1 ps.

#### V. Reconstruction of free energy surface (FES) from the RAM simulations

After about 300 ns, the sampling along the first 3 CVs reached convergence with fluctuations of bias within 1 kcal/mol, allowing a reliable reconstruction of the corresponding FES. The production run was continued for another 100 ns to sample enough conformations. These conformations were first clustered into microstates using the regular spatial method (cut-off radius of 0.13), and then the free energy of each state was reweighted according to the deposited potential along each CV, which can be obtained from the analysis module of METAGUI.^80^ To visualize the distribution of these microstates and their relative energy differences, we further performed principal component analysis to project the microstates represented in the 3-dimensional CV space into 2-dimensional space spanned by PC1 and PC2. Then we plotted these microstates in a 3-dimensional space spanned by PC1, PC2, and free energy differences *ΔG*.

#### VI. Independent validation of chemical shifts with SPARTA+

During the REX and RAM simulations, the chemical shifts were computed via CamShift in ALMOST 2.1.^78^ As an independent validation for the efficacy of the bias, we further calibrated the chemical shifts of 2000 snapshots with MDTraj^81^ and SPARTA+.^34^

#### VII. Adaptive sampling

The first round of adaptive sampling started from the snapshots of low-energy microstates obtained from the previous step, *i.e.*, 1200 structures for the apo form, 400 structures for the binary form, and 200 structures for the ternary form. The initial velocities were randomly generated to satisfy the Maxwell distribution at 300K. For the apo form, a 10 ns simulation was performed for each run, whereas for the binary, each simulation lasted 30 ns, resulting in a total of 12 μs trajectories for both the apo and binary forms. Three rounds of adaptive sampling were started from the 400 microstates, derived via K-mean clustering of all snapshots of previous ensembles, to obtain a converged free energy landscape. Therefore, a total of 100 µs trajectories and 100,0000 snapshots (100 ps per frame) were collected for both the apo and binary form after three rounds of adaptive sampling, and a total of 60 µs trajectories were collected for the ternary form.

#### VIII. Markov state model (MSM) and time-lagged independent component analysis (tICA)

The Cartesian coordinates of key hydrophobic residues, including R-spine residues, L95, L106, Y164, and F185, and the shell residues, V104, M118, and M120, were chosen as the metrics to characterize the conformational transition of the hydrophobic core of PKA-C. Specifically, each snapshot was first aligned to the same reference structure by superimposing the αE (residues 140-160) and αF helices (residues 217-233) and represented by the deviation of the Cartesian coordinates of key residues. The representation in this metric space was further reduced to 10-dimension vectors using time-lagged independent component analysis (tICA)^82^ at a lag time of 1 ns. All snapshots were clustered into 400 microstates with K-mean clustering. An MSM was built upon the transition counts between these microstates.

#### IX. Kinetic Monte Carlo trajectory of PKA-C in different forms

Long trajectories were generated using a kinetic Monte Carlo method based on the MSM transition probability matrix of the three forms of PKA-C. Specifically, the discrete jumps between the 100 microstates were sampled for 60 μs. Random conformations were chosen for each state from all the available snapshots. Subsequently, the time series of various order parameters were analyzed and plotted.

### Protein expression and purification

The recombinant human Cα subunit of cAMP-dependent protein kinase with the Phe to Ala mutation in position 100 (PKA-C^F100A^) was generated from the human PKA-Cα wild-type using Quik-Change Lightning mutagenesis kit (Agilent genomics). The key resource table lists the PCR primers used to modify the pET-28a expression vector encoding for the wild-type human PKA-Cα gene (*PRKACA –* uniprot P17612) ^21–23^. The unlabeled and uniformly ^15^N-labeled PKA-C^F100A^ mutant was expressed and purified following the same protocols used for the wild-type protein.^41^ Briefly, transformed *E. coli* BL21 (DE3) *pLyss* cells (Agilent) were cultured overnight at 30 °C in Luria-Bertani (LB) medium. The next morning, the cells were transferred to fresh LB medium for the overexpression of the unlabeled protein or to M9 minimal medium supplied with ^15^NH_4_Cl (Cambridge Isotope Laboratories Inc.) as the only nitrogen source for the labeled protein overexpression. In both cases, protein overexpression was induced with 0.4 mM of **β**-D-thiogalactopyranoside (IPTG) and carried out for 16 hours at 20 °C. The cells were harvested by centrifugation and resuspended in 50 mM Tris-HCl, 30 mM KH_2_PO_4_, 100 mM NaCl, 5 mM 2-mercaptoethanol, 0.15 mg/mL lysozyme, 200 μM ATP, DNaseI, 1 tablet of cOmplete ULTRA EDTA-free protease inhibitors (Roche Applied Science) (pH 8.0) and lysed using French press at 1,000 psi. The cell lysate was cleared by centrifugation (60,000 × g, 4 °C, 45 min), and the supernatant was batch-bound with Ni^2+^-NTA agarose affinity resin (Thermo Scientific). The his-tagged PKA-C^F100A^ was eluted with 50 mM Tris-HCl, 30 mM KH_2_PO_4_, 100 mM NaCl, 5 mM 2-mercaptoethanol, 0.5 mM phenylmethylsulfonyl fluoride (PMSF) (pH 8.0) supplied with 200 mM of imidazole. The tail of poly-His was cleaved using a stoichiometric amount of recombinant tobacco etch virus (TEV) protease in 20 mM KH_2_PO_4_, 25 mM KCl, 5 mM 2-mercaptoethanol, 0.1 mM PMSF (pH 6.5), overnight at 4 °C. The different phosphorylation states of PKA-C^F100A^ were separated using a cation exchange column (HiTrap Q-SP, GE Healthcare Life Sciences) using a linear gradient of KCl in 20 mM KH_2_PO_4_ at pH6.5.^83^ The purified protein isoforms were then stored in phosphate buffer containing 10 mM dithiothreitol (DTT), 10 mM MgCl_2_, and 1 mM NaN_3_ at 4 °C. The protein purity was assessed by sodium dodecyl sulfate-polyacrylamide gel electrophoresis (SDS–PAGE).

### Peptide synthesis

The Kemptide (LRRASLG) and PKI_5–24_ (TTYADFIASGRTGRRNAIHD) peptides were synthesized using a CEM Liberty Blue microwave synthesizer using standard Fmoc chemistry. All peptides were cleaved with Reagent K (82.5% TFA, 5% phenol, 5% thioanisole, 2.5% ethanedithiol, and 5% water) for 3 h and purified using a semipreparative Supelco C18 reverse-phase HPLC column at 3 mL/min. The purified peptides were concentrated, lyophilized, and stored at −20 °C. Molecular weight and quantity were verified by MALDI-TOF and/or amino-acid analysis (Texas A&M University).

### Isotherm titration calorimetry (ITC) measurements

PKA-C^F100A^ was dialyzed into 20 mM MOPS, 90 mM KCl, 10 mM DTT, 10 mM MgCl_2_, and 1 mM NaN_3_ (pH 6.5) and concentrated using conical spin concentrator (10 kDa membrane cut-off, Millipore) to a solution at 80-100 μM, as confirmed by A280 = 55,475 M^−1^ cm^−1^. Approximately 300 μL of protein was used for each experiment, with 50 μL of 2 mM ATPγN and/or 1 mM PKI_5–24_ in the titrant syringe. All measurements were performed at 300 K in triplicates with a low-volume NanoITC (TA Instruments). The binding was assumed to be 1:1, and curves were analyzed with the NanoAnalyze software (TA Instruments) using the Wiseman isotherm ^49^

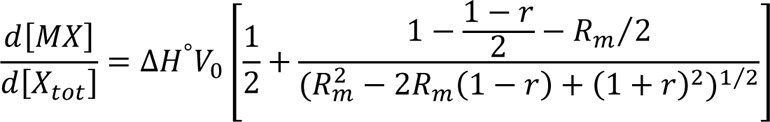

where *d*[*MX*] is the change in total complex relative to the change in total protein concentration, *d*[*X_tot_*] is dependent on *r* (the ratio of *K*_d_ relative to the total protein concentration), and *R_m_* (the ratio between total ligand and total protein concentration). The heat of dilution of the ligand into the buffer was considered for all experiments and subtracted.

The free energy of binding was determined from:

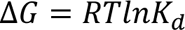

where *R* is the universal gas constant and *T* is the temperature at measurement (300 K). The entropic contribution to binding was calculated using:

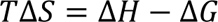

The degree of cooperativity (σ) was calculated as:

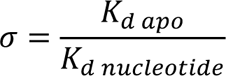

where *K_d apo_* is the dissociation constant of PKI_5–24_ binding to the apo-enzyme, and *K_d Nucleotide_* is the corresponding dissociation constant for PKI_5–24_ binding to the nucleotide-bound kinase.

### Enzyme assays

Steady-state coupled enzyme activity assays using Kemptide as substrate were performed under saturating ATP concentrations and spectrophotometrically at 298 K, as reported by Cook *et al.*^47^ The values of *V_max_* and *K_M_* were obtained from a nonlinear fit of the initial velocities to the Michaelis-Menten equation.

### NMR spectroscopy

NMR measurements were performed on a Bruker Avance NEO spectrometer operating at a ^1^H Larmor frequency of 600 MHz equipped with a cryogenic probe or on a Bruker Avance III 850 MHz spectrometer equipped with a TCI cryoprobe. The NMR experiments were recorded at 300K in 20 mM KH_2_PO_4_ (pH 6.5), 90 mM KCl, 10 mM MgCl_2_, 10 mM DTT, 1 mM NaN_3_, 5% D_2_O, and 0.1% 4-benzene sulfonyl fluoride hydrochloride (AEBSF, Pefabl–c - Sigma-Aldrich). Concentrations for samples were 0.15 mM of uniformly ^15^N-labeled PKA-C^F100A^, as determined by A_280_ measurements, 12 mM ATPγN or ADP was added for the nucleotide-bound form, and 0.3 mM PKI_5-24_ for the ternary complex. [^1^H, ^15^N]-WADE-TROSY-HSQC pulse sequence^84^ was used to record the amide fingerprint spectra of PKA-C^F100A^ in the apo, nucleotide-bound (ADP- or ATPγN-bound – binary form), and ternary complex (PKAC^F100A^/ATPγN/PKI_5-24_). All [^1^H, ^15^N]-WADE-TROSY-HSQC experiments were acquired with 2048 (proton) and128 (nitrogen) complex points, processed using NMRPipe^85^ and visualized using NMRFAM-SPARKY ^86^ and POKY.^87^ Combined chemical shift perturbations (CSP) were calculated using ^1^H and ^15^N chemical shifts according to:

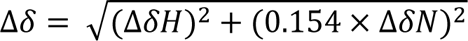

in which Δδ is the CSP; Δδ*_H_* and Δδ*_N_* are the differences of ^1^H and ^15^N chemical shifts, respectively, between the first and last point of the titration; and 0.154 is the scaling factor for nitrogen.^88^ ***COordiNated ChemIcal Shift bEhavior (CONCISE).*** The normal distributions reported in the CONCISE plot were calculated using principal component analysis for residues whose chemical shifts responded linearly to ligand binding.^51^ In this work, we use the ^1^H and ^15^N chemical shifts derived from the [^1^H, ^15^N]-WADE-TROSY-HSQC experiments for the apo, ADP, ATPγN, and ATPγN/PKI_5–24_ bound forms of PKA-C.

### CHEmical Shift Covariance Analysis (CHESCA)

This analysis was used to identify and characterize allosteric networks of residues showing concerted responses to nucleotide and pseudosubstrate binding. To identify inter-residue correlations, four states were used: apo, ATPγN-bound, ADP-bound, and ATPγN/PKI_5–24_. The identification of inter-residue correlations by CHESCA relies on agglomerative clustering (AC) and singular value decomposition (SVD).^53^ Pairwise correlations of chemical shift changes were calculated to identify networks. When plotted on a correlation matrix, the 2D correlations allow the identification of local and long-range correlated residues. For each residue, the maximum change in the chemical shift was calculated in both the ^1^H (*x*) and ^15^N (*y*) dimensions (Δδ*_x,y_*). The residues included in the CHESCA analysis were those satisfying Δδ_x,y_ > ½ *Δν*_x*A*,y*A*_+½ *Δν*_x*B*,y*B*_, where A and B correspond to two different forms analyzed, and Δν denotes the linewidth. Correlation scores were used to quantify the CHESCA correlation of a single residue or a group of residues with another group. Correlation scores were evaluated for single residues or for the entire protein using the following expression:

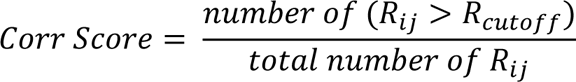

where *R_ij_* is the correlation coefficient and *i* and *j* denote (a) a single residue and the remainder residues of the protein, respectively, or (b) both represent all the assigned residues in the entire protein. For all the analyses a *R_cutoff_* of 0.98 was used. Community CHESCA analysis utilizes a similar approach to map correlations between functional communities within the kinase. Each community represents a group of residues associated with a function or regulatory mechanism.^55^ To represent community-based CHESCA analysis, we utilized *R_cutoff_* > 0.8. For instance, to represent a chemical shift correlation between two community (X and Y) with *n_A_* and *n_B_* number of assigned residues, respectively, the correlation score between A and B is defined as

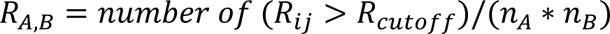

where *R_ij_* is the correlation coefficient between residue *i* (belonging to community A) and residue *j* (belonging to community B). *R_cutoff_* is the correlation value cutoff. *R_A,B_* assumes values from 0 (no correlation between residues in A and B) to 1 (all residues in A have correlations > cut-off with all residues in B).

## Acknowledgments

This work was supported by the National Institute of Health GM 100310 and HL 144130 to GV. The authors would like to acknowledge the Minnesota Supercomputing Institute for MD calculations. YW would like to thank the Guangdong Pearl River Talent Program (2021QN02Y618) and the National Natural Science Foundation of China (22007069, 92269102) for part of the MD analysis carried out at the Shenzhen Bay Laboratory Supercomputing Centre.

## Competing interest

The authors declare no financial and non-financial competing interests.

## Author contributions

Y.W., C.O., and G.V. designed research. Y.W. performed the MD simulations and analysis. C.O. prepared the kinase samples, executed all NMR and ITC experiments, and analyzed the results. C.W. performed the kinase activity assay and prepared the peptides for the assay. C.O., Y.W. contributed to the manuscript, preparing all the text and figures. M.V.S., K.N.H., C.C., J.G., and M.V. contributed to the critical analysis of the data and writing of the manuscript. D.A.B. contributed to the critical analysis of the data and writing the manuscript. S.S.T. contributed to the writing of the manuscript. G.V. designed and directed the experiments and contributed to the writing of the manuscript. All authors have approved the final version of the manuscript.

To whom correspondence should be addressed.

## Data Availability

All the data generated or analyzed in this study are included in the manuscript and supporting files. The NMR chemical shifts and MD trajectories are deposited in the Data Repository for the University of Minnesota, https://hdl.handle.net/11299/261043.

---

**Figure 2 – figure supplement 1.**
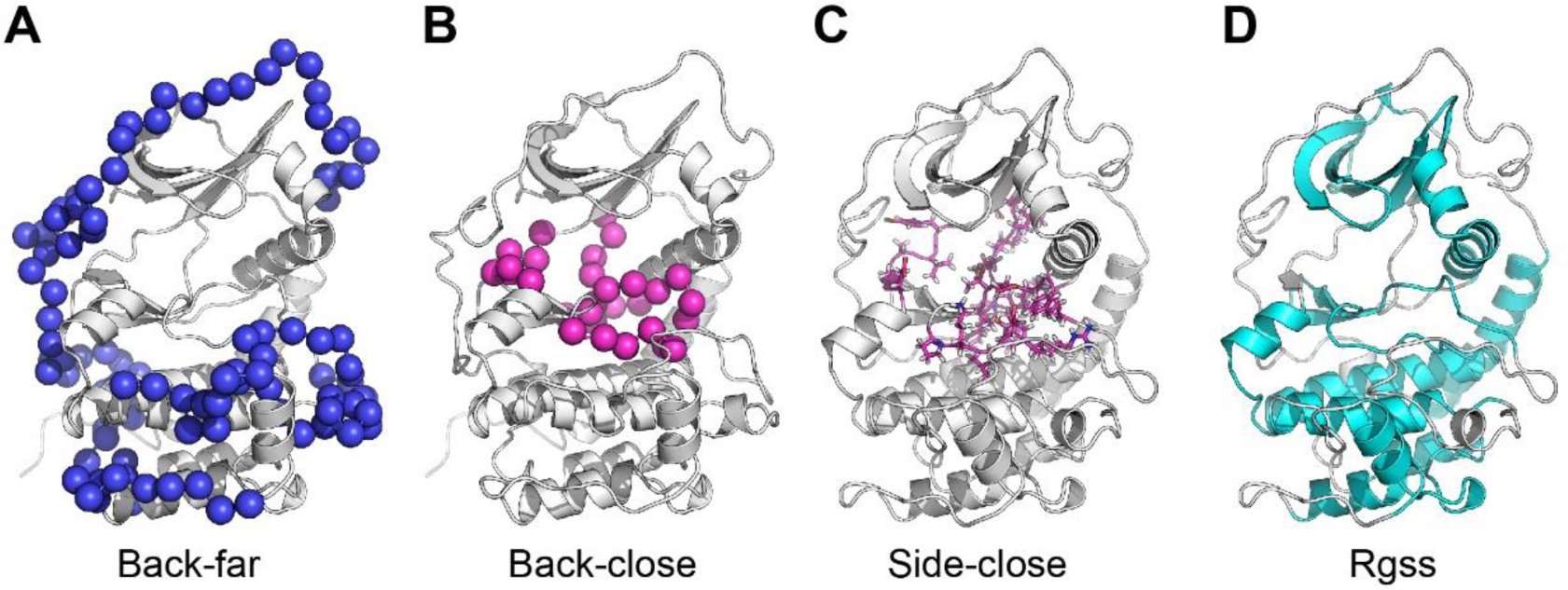
Illustration of the collective variables (CVs) used in the RAM simulations. (**A**) Backbone ψ angles of loops not in contact with ATP (Back-far). The Cα atoms are depicted as blue spheres. (**B**) Backbone ψ angles of loops in contact with ATP (Back-close). The Cα atoms are colored in magenta. (**C**) Side chains χ_1_ angles of loops in contact with ATP (Side-close). The side chains are represented in sticks colored in magenta. (**D**) The radius of gyration is calculated over the rigid part of the protein (Rgss), where the residues involved are colored in cyan.

**Figure 2 – figure supplement 2.**
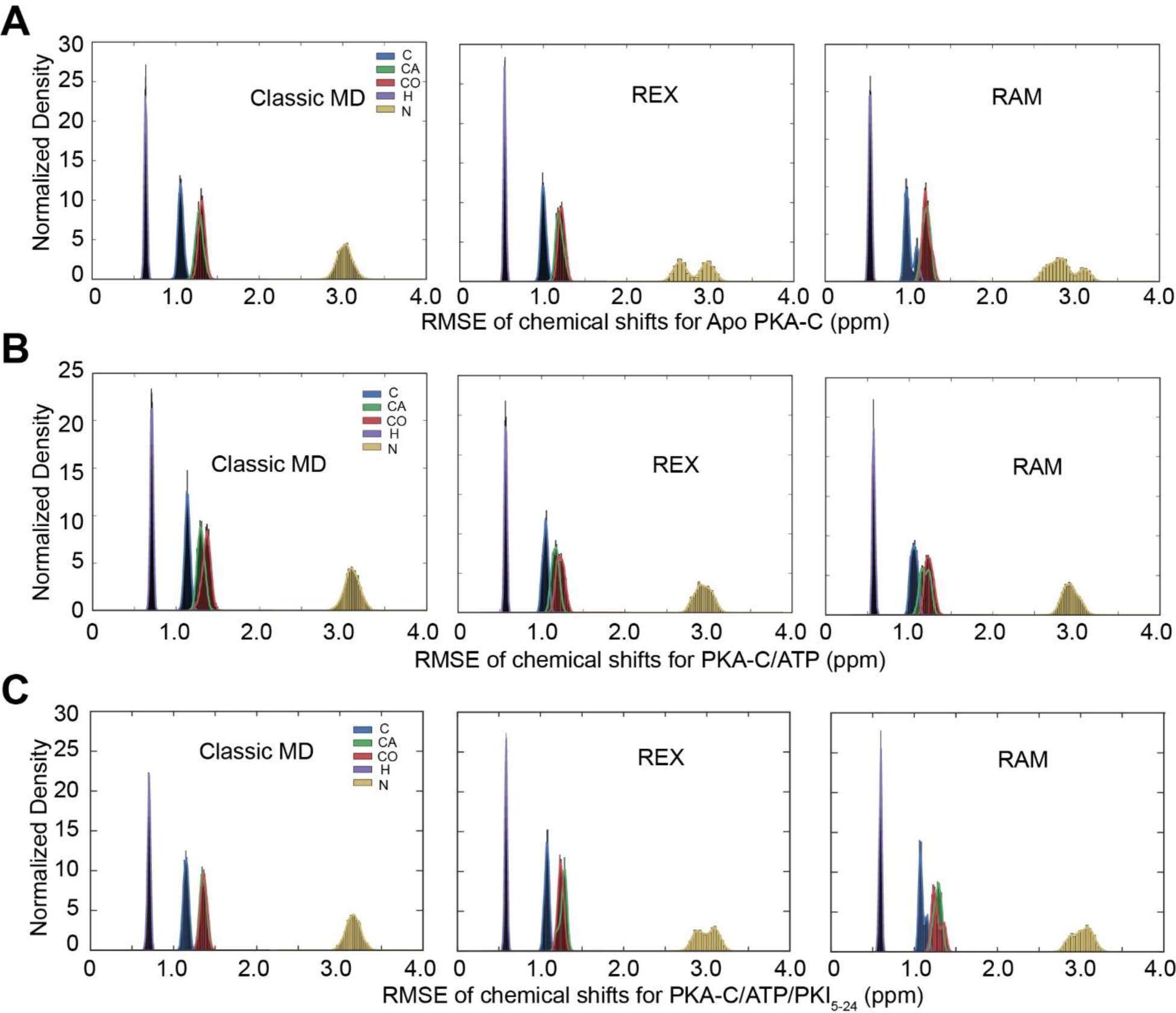
Distribution of the Root-Mean-Square-Error (RMSE) of the chemical shifts for the different simulation schemes. (**A**) RMSE of CS for apo PKA-C calculated from standard MD (left), REX (middle), and RAM (right). (**B**) RMSE of CS for PKA-C/ATP calculated from standard MD (left), REX (middle), and RAM (right). (**C**) RMSE of CS for PKA-C/ATP/PKI_5-24_ from standard MD (left), REX (middle), and RAM (right). Color codes for different backbone atoms (C, Cα, CO, H, and N) are indicated in the left figures.

**Figure 2 – figure supplement 3.**
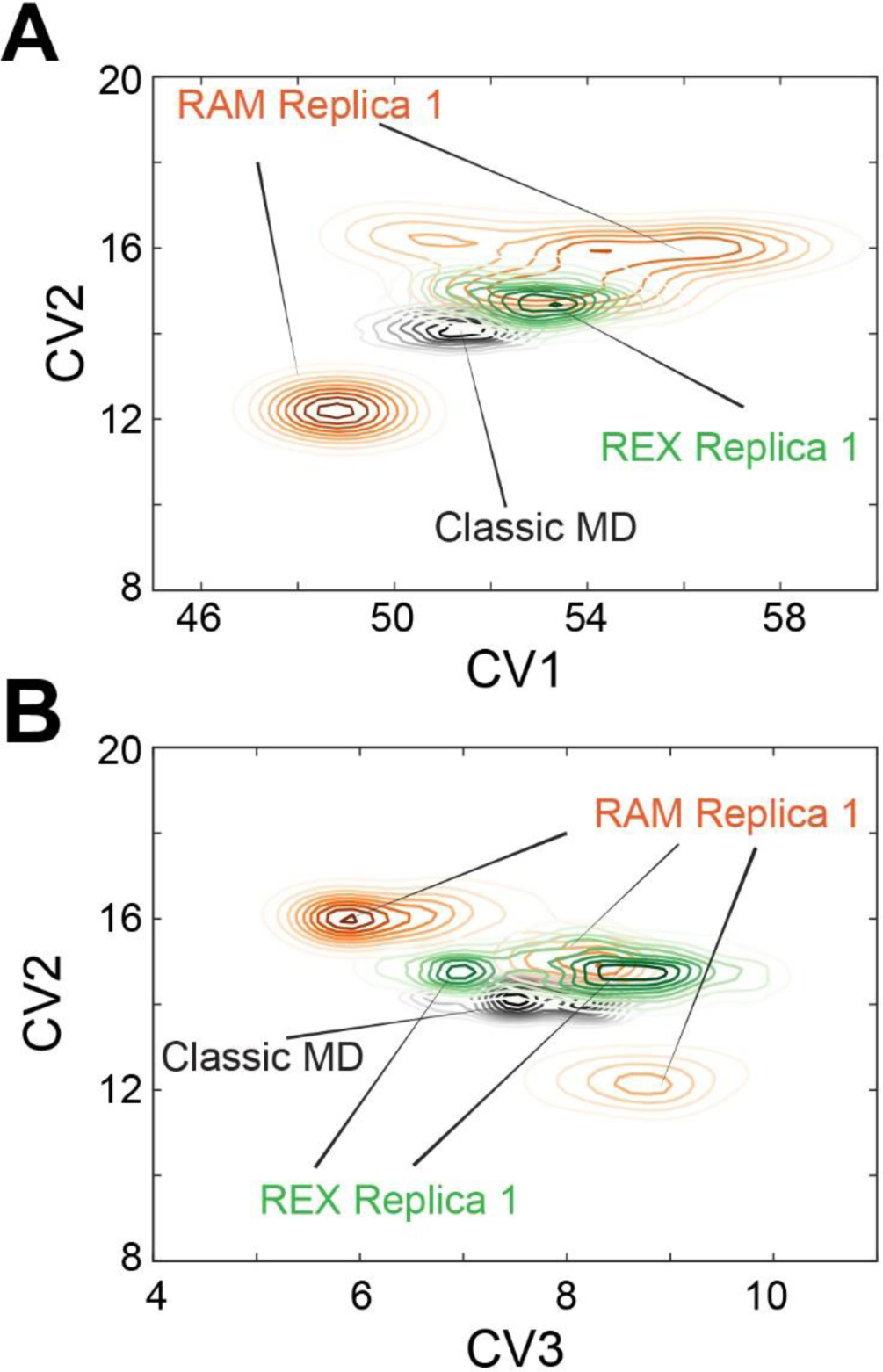
Replica-averaged metadynamics (RAM) simulations explore a larger conformational space than standard MD and replica exchange (REX) simulations. (**A**) Comparison of conformational space (CV1 vs. CV2) sampled by RAM Replica 1, standard MD, and REX Replica 1 for PKA-C. (**B**) Comparison of conformational space (CV3 vs. CV2) sampled by RAM Replica 1, standard MD, and REX Replica 1 for apo PKA-C.

**Figure 2 – figure supplement 4.**
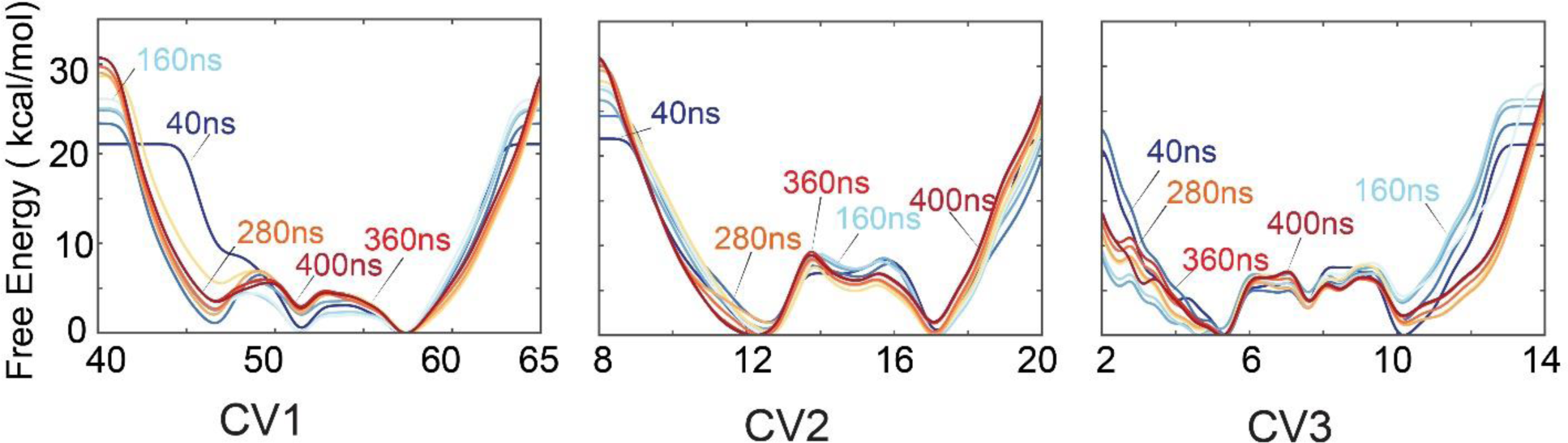
Accumulative deposition of history-dependent biases along the first three CVs for the RAM simulations of the apo PKA-C. The accumulative biases converged after ∼300 ns in the three CVs.

**Figure 3 – figure supplement 1.**
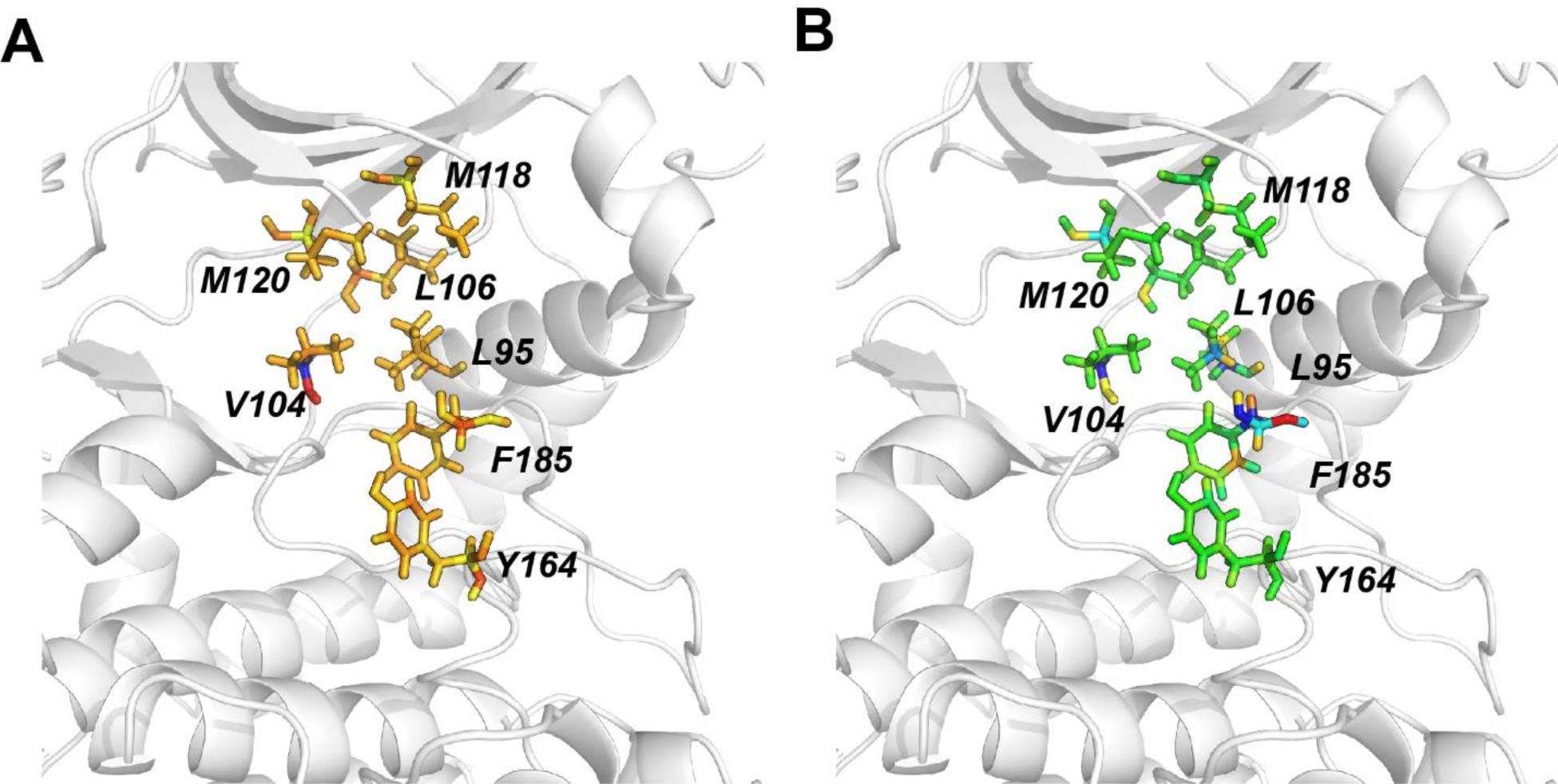
R spine and shell residues selected for two time-lagged independent components (tICA) and Markov State Model (MSM) analysis. (**A**) Atom motions of key residues that define tIC1 of apo PKA-C colored according to the superposition deviations. Backbone atoms of Val104 show the largest change in tIC1. (**B**) Atom motions of key residues that define tIC2 of apo PKA-C colored according to the superposition deviations. Backbone atoms of Phe185 and Val104 show the largest change in tIC2.

**Figure 3 – figure supplement 2.**
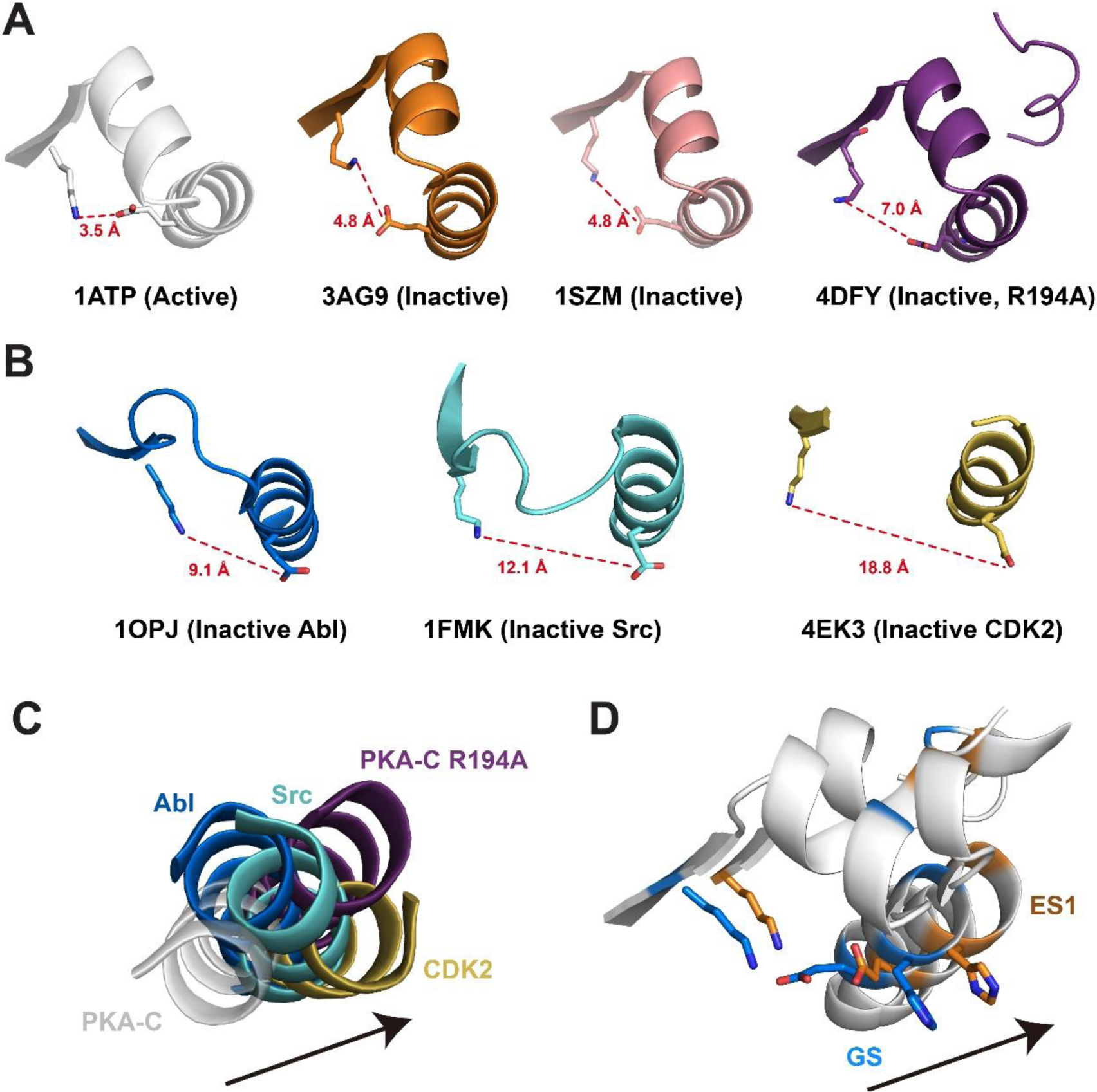
Structural features of αC-out transition (ES1) in various inactive kinases. (**A**) Crystal structures of PKA-C in active and inactive states highlight the αB-αC loop alterations. Crystal structures of PKA-C in the active (1ATP) and inactive conformations (3AG9, 1SZM, 4DFY), highlighting the electrostatic interactions between K72 in β3 and E91 in theαC. (**B**) Disruption of the K72 - E91salt bridge in the inactive structures of Abl (1OPJ), Src (1FMK), and CDK2 (4EK3). (**C**) The αC helix orientation of active PKA-C compared to the orientation in inactive kinases. (**D**) Structural transition from the GS state to ES1 state characterized by the outward movement of the αC helix (i.e., kinase inactivation).

**Figure 3 – figure supplement 3.**
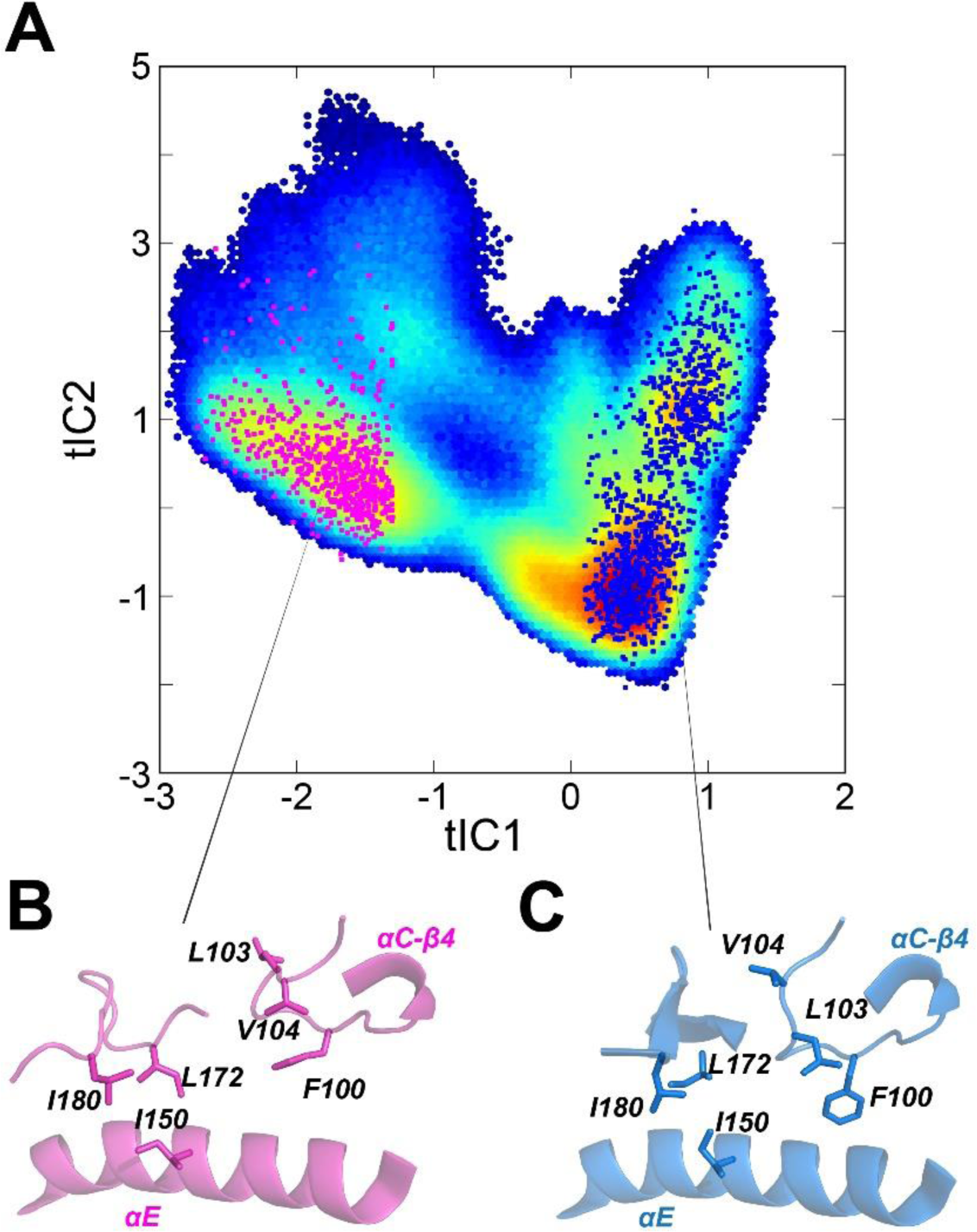
Distinct hydrophobic packing for residues around the αC-β4 loop in the GS and ES states of apo PKA-C. (**A**) Projections of randomly selected conformations for the GS (blue) and ES (magenta) onto the conformational landscape of apo PKA-C. Snapshots with tIC1 < 1.2 were clustered to separate the ES and GS, whereas those confomers with tIC1 > 0.2 were clustered as GS. (**B,C**) Close up of the hydrophobic packing in the ES and GS (**C**) states, highlighting Leu103, Val104, Ile150, Leu172, and Ile180 that show slow exchange in the CPMG experiments.

**Figure 5 - figure supplement 1.**
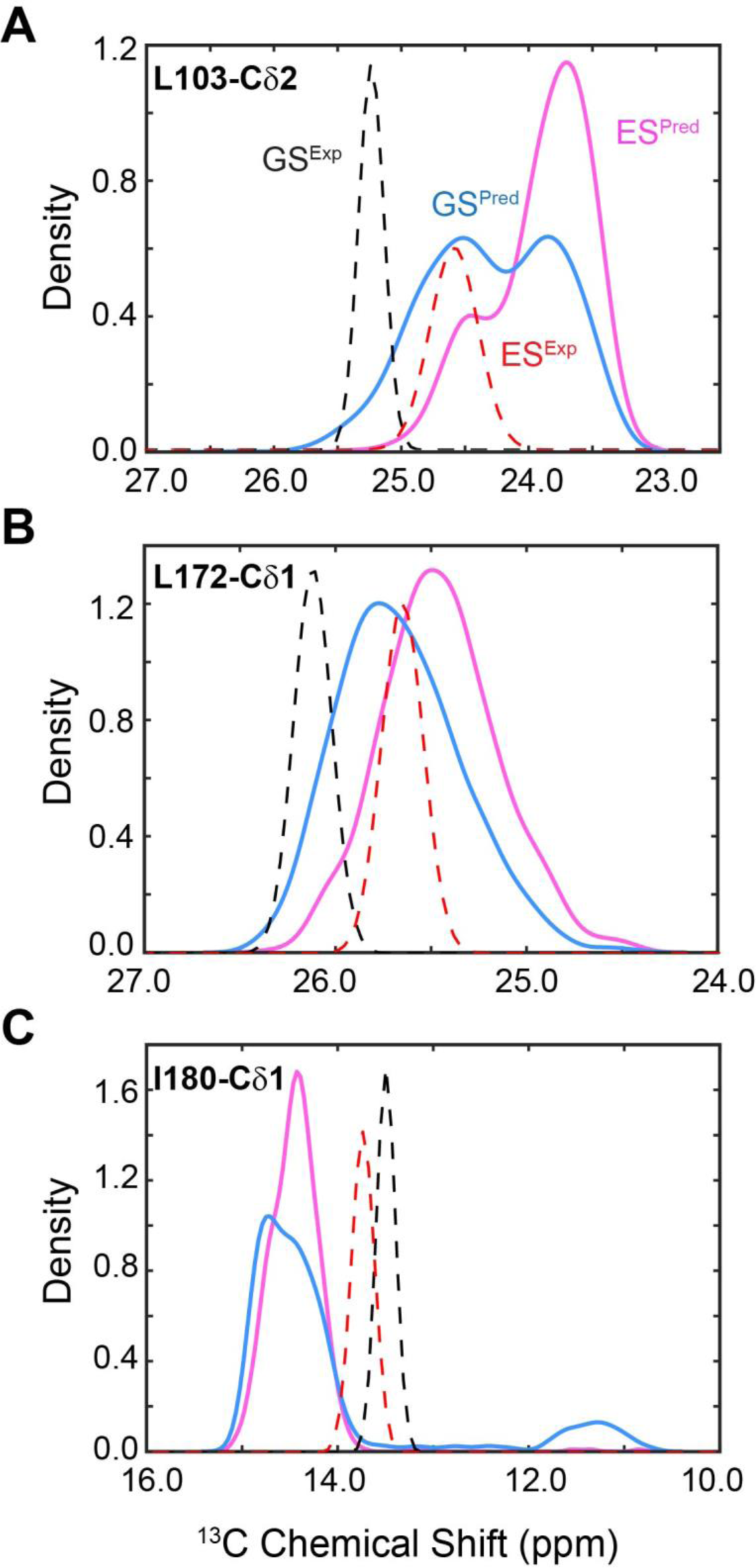
Distribution of predicted and experimental ^13^C CS of selected methyl groups. **(A-C)** ES (magenta) and GS (blue) of apo PKA-C for Leu103-Cδ2 (**A**), Leu172-Cδ1 (**B**), and Ile180-Cδ1 (**C**). The experimental CSs are shown as dotted lines for GS (black) and ES (red).

**Figure 6-figure supplement 1.**
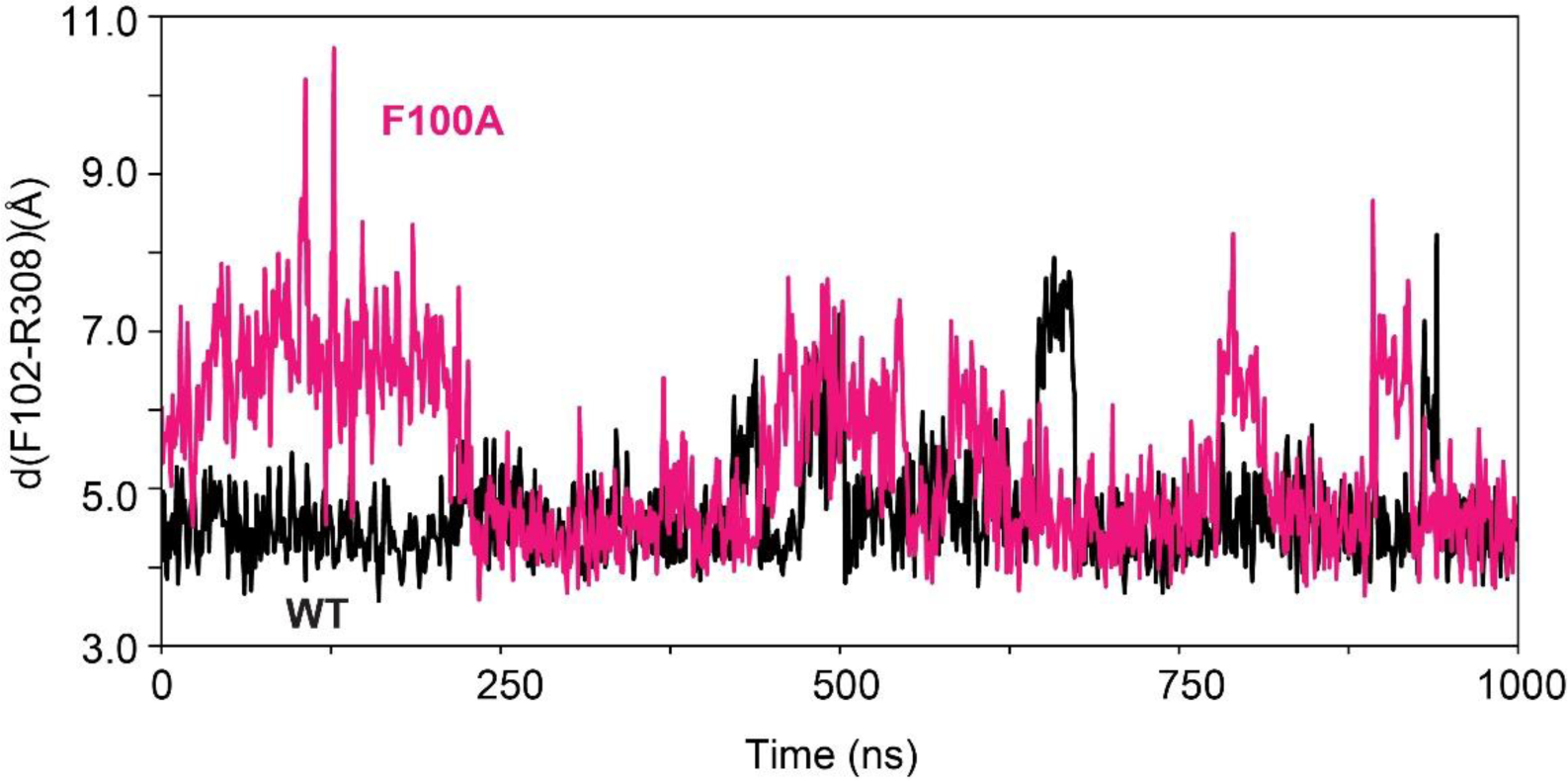
Time series of the distance between F102 and R308 for PKA-C^WT^ (black) and the PKA-C^F100A^ mutant (magenta). The π-cation interactions between the aromatic ring of F102 and the guanidine group of R308 are more persistent in the WT enzyme than in the F100A mutant.

**Figure 9 - figure supplement 1.**
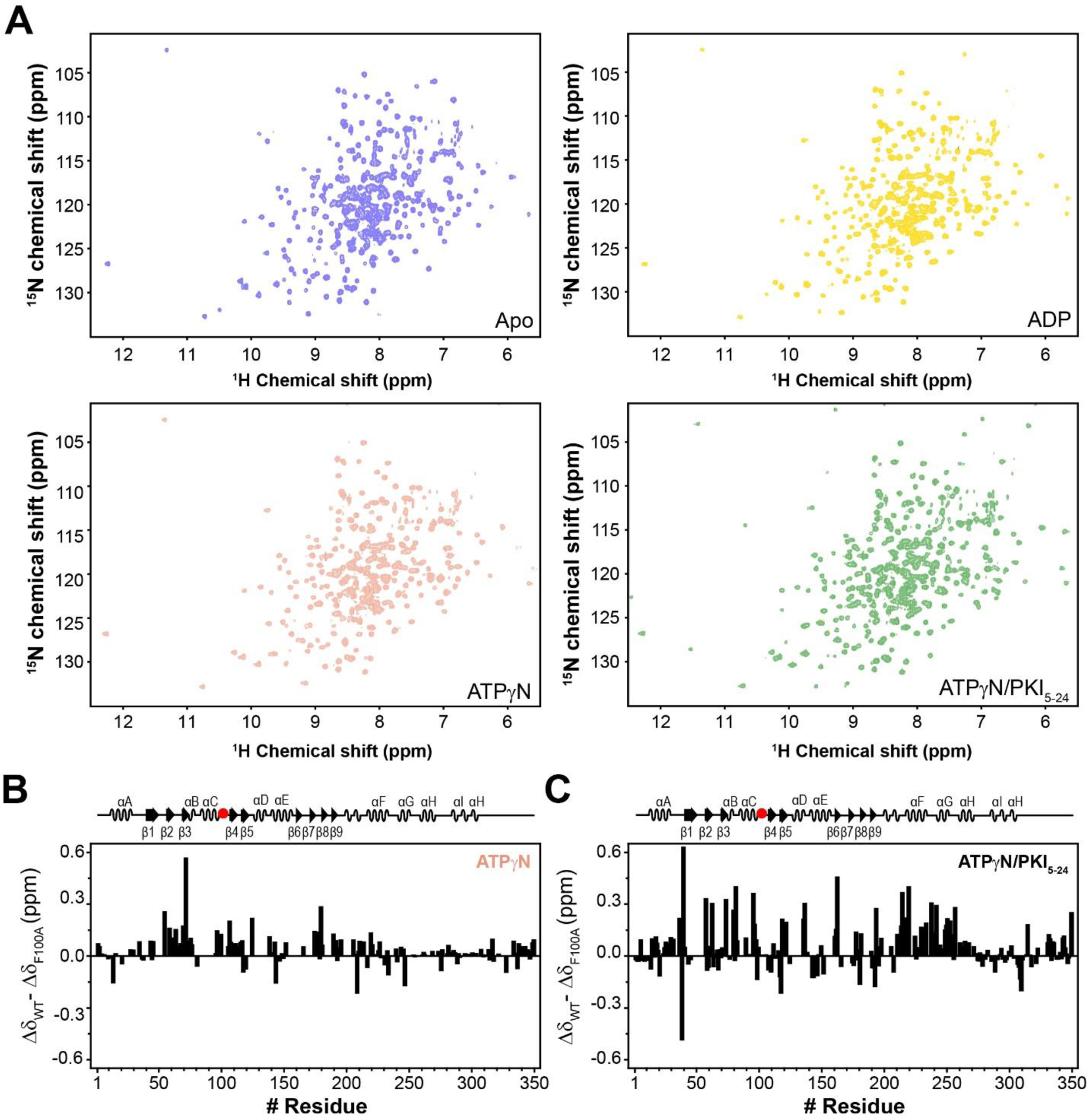
NMR fingerprints of PKA-C^F100A^. (**A**) [^1^H,^15^N]-WADE-TROSY spectra of apo, ADP-bound, ATPγN-bound, and ATPγN/PKI_5-24_ bound PKA-C^F100A^. (**B**) Changes in the chemical shift perturbation (CSP) between PKA-C^WT^ and PKA-C^F100A^ bound to ATPγN. (**C**) Changes in CSP (Δδ_WT_ - Δδ_F100A_) upon binding ATPγN and PKI_5-24_.

**Figure 9 - figure supplement 2.**
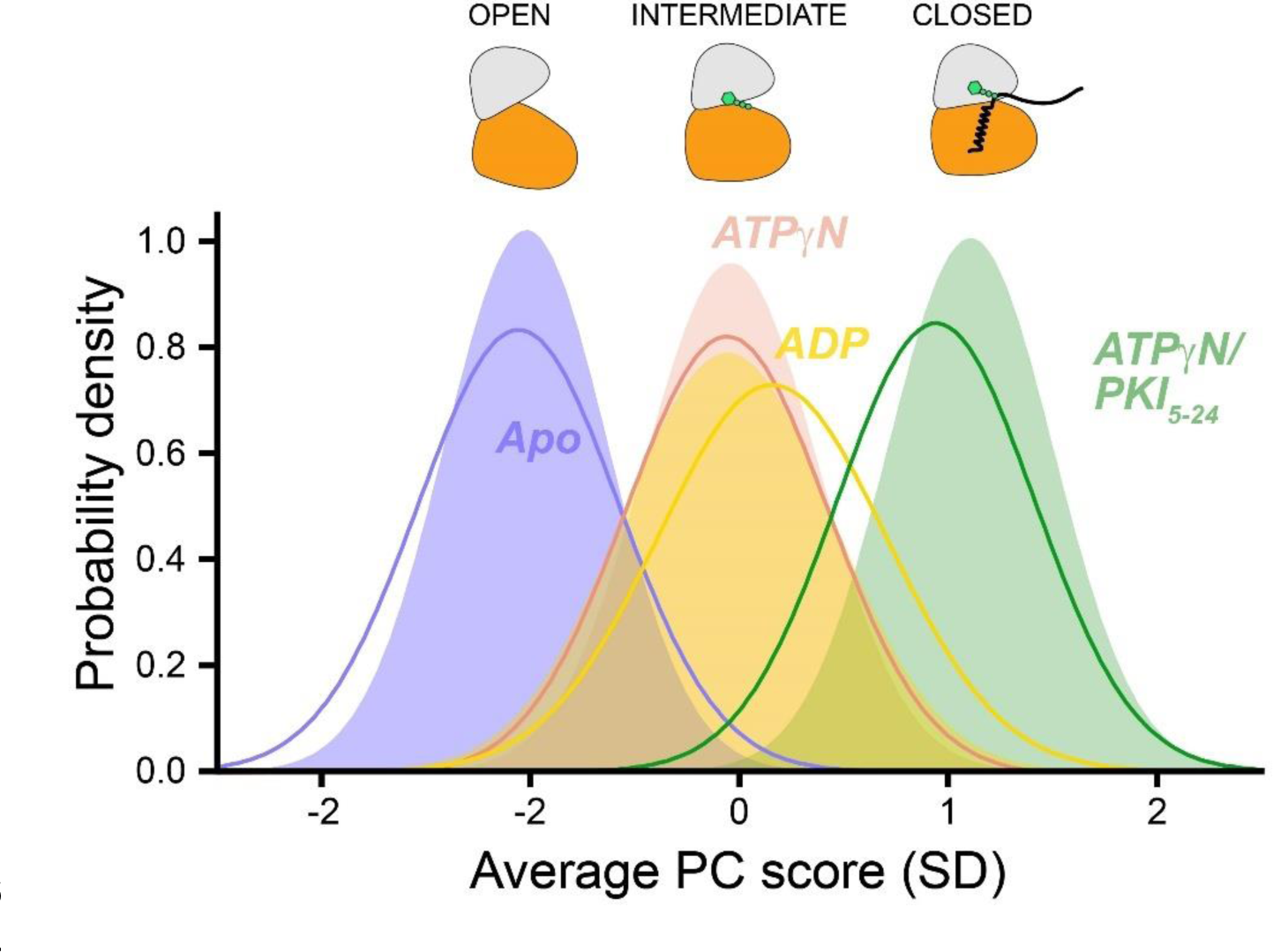
CONCISE plot showing the shifts of the probability distribution of the amide resonances as a function of nucleotides and substrate binding. The per-residue chemical shift information is averaged into the average principal component (PC) score indicative of the position of each conformational state of the kinase along the open-to-closed equilibrium.

**Figure S10 - figure supplement 2.**
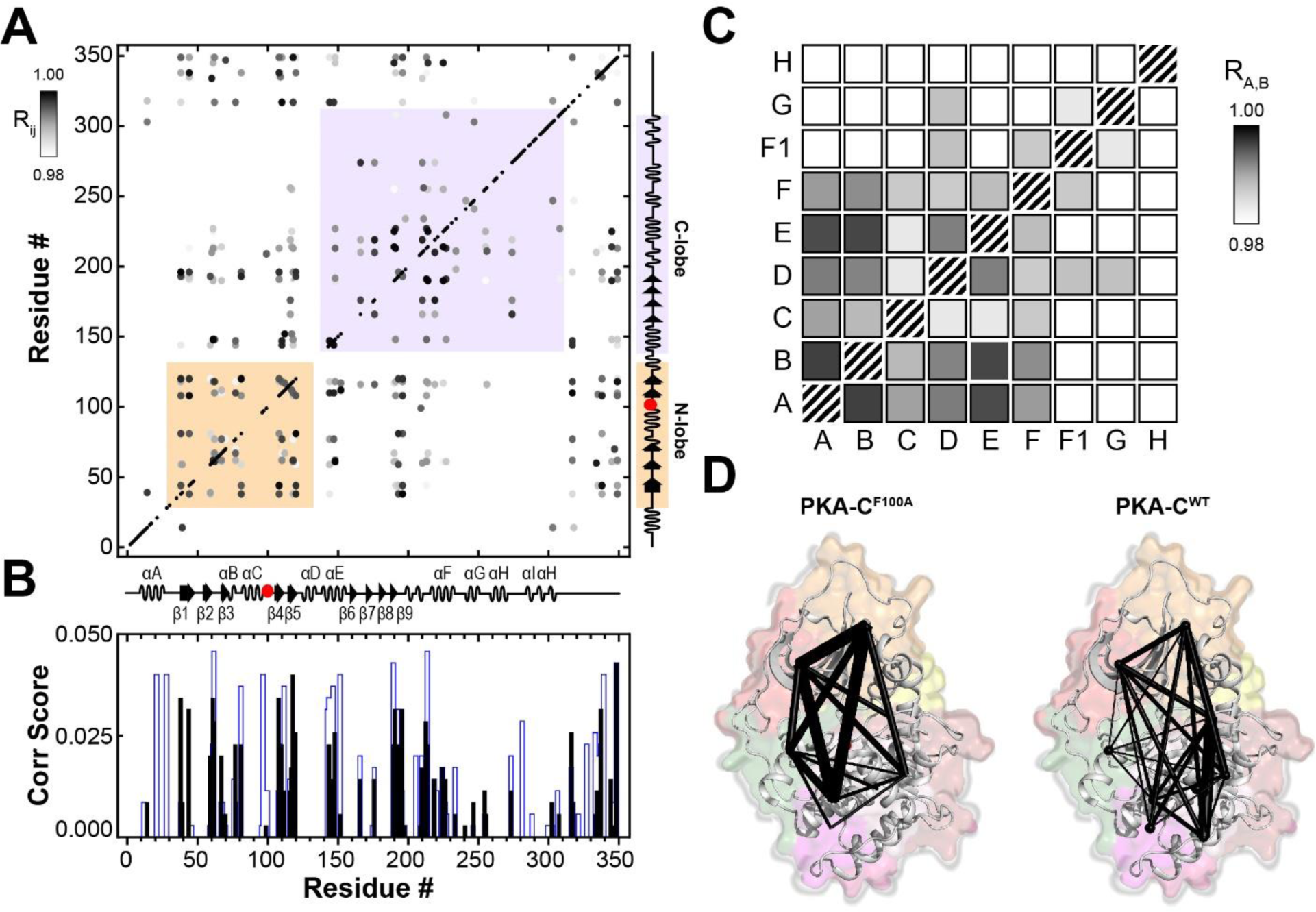
Intermolecular allosteric network of F100A mapped using CHESCA and community CHESCA. (**A**) CHESCA matrix obtained from the amide chemical shift trajectories of PKA-C^F100A^ in the apo, ADP-bound, ATPγN-bound, and ATPγN/PKI_5-24_-bound states. Only correlations with R_ij_ > 0.98 are displayed. (**B**) Plot of the correlation scores vs. residue calculated for PKA-C^WT^ (blue) and PKA-C^F100A^ (black). (**C**) Community CHESCA analysis of PKA-C^F100A^. Only correlations with R_A,B_ > 0.98 are shown. (**D**) Spider plots indicating the correlated structural communities of PKA-C^F100A^ and PKA-C^WT^ plotted on their corresponding structures. The size of each node is independent of the number of residues it encompasses, and the weight of each line indicates the strength of coupling between the individual communities.

**Figure 2 – supplementary Table 1.** ΔG (kcal/mol) and relative population of the ground state and the first 6 excited states in different forms of PKA-C obtained from the RAM simulations.

|  | GS | ES1 | ES2 | ES3 | ES4 | ES5 | ES6 |
| --- | --- | --- | --- | --- | --- | --- | --- |
| Apo | 0<br>(58.0%) | 0.38<br>(30.8%) | 1.22<br>(7.6%) | 2.10<br>(1.8.%) | 2.28<br>(1.3%) | 2.85<br>(0.5%) | 8.16*<br>($<1e-4$) |
| Binary | 0<br>(99.4%) | 3.11<br>(0.6%) | 5.80<br>($<1e-4$) | 6.66<br>($<1e-4$) | 7.25<br>($<1e-4$) | 8.27<br>($<1e-4$) | 8.47<br>($<1e-4$) |
| Ternary | 0<br>(100.0%) | 4.85<br>($<1e-4$) | 5.68<br>($<1e-4$) | 6.92<br>($<1e-4$) | 7.04<br>($<1e-4$) | 7.58<br>($<1e-4$) | 7.87<br>($<1e-4$) |

\* Numbers in red refer to populations of excited states below 0.5%.

**Supplementary Table 2.** Kinetic parameters of Kemptide phosphorylation by PKA-C^WT^ and PKA-C^F100A^ obtained from coupled assays. The *K_M_* and *V_max_* values were obtained from a non-linear least squares analysis of the concentration-dependent initial phosphorylation rates. Errors in the *k_cat_*/*K_M_* ratios were propagated from the individual errors in *K_M_* and *k_cat_*.

|  | **PKA-C<sup>WT</sup>** | **PKA-C<sup>F100A</sup>** |
| --- | --- | --- |
| $V_{max}$ | $0.322 \pm 0.005$ | $0.379 \pm 0.009$ |
| $K_M$ | $30 \pm 1$ | $42 \pm 3$ |
| $k_{cat}$ | $15 \pm 1$ | $17 \pm 1$ |
| $k_{cat}/K_M$ | $0.50 \pm 0.04$ | $0.41 \pm 0.08$ |

**Supplementary Table 3.** Changes in enthalpy, entropy, free energy, and dissociation constants for nucleotide binding to PKA-C^WT^ and PKA-C^F100A^. All errors were calculated from triplicate measurements. Values for PKA-C^WT^ are taken from Walker *et al.* ^22^

| | $K_d$ (μM) | $\Delta G$ (kcal/mol) | $\Delta H$ (kcal/mol) | $-T\Delta S$ (kcal/mol) | $\sigma$ |
| --- | --- | --- | --- | --- | --- |
| PKA-C<sup>WT</sup> | 83 ± 8 | -5.61 ± 0.06 | -3.6 ± 0.1 | -2.0 ± 0.1 | N/A |
| PKA-C<sup>F100A</sup> | 73 ± 2 | -5.7 ± 0.2 | -21 ± 5 | 7 ± 3 | N/A |

**Supplementary Table 4.** Changes in enthalpy, entropy, free energy, and dissociation constants for PKI_5-24_ binding to apo and ATPγN - saturated PKA-C^WT^ and PKA-C^F100A^. All errors were derived from triplicate measurements. The error for the cooperativity coefficient (σ) was propagated from the errors in *K*_d_. Values for PKA-C^WT^ were originally published in Walker *et al.* ^22^.

| PKI<sub>5-24</sub> to apo kinase |  |  |  |  |  |
| --- | --- | --- | --- | --- | --- |
| | $K_d$ (μM) | $\Delta G$ (kcal/mol) | $\Delta H$ (kcal/mol) | $-T\Delta S$ (kcal/mol) | $\sigma$ |
| PKA-C<sup>WT</sup> | $17 \pm 2$ | $-6.57 \pm 0.08$ | $-10.8 \pm 0.5$ | $4.2 \pm 0.5$ | N/A |
| PKA-C<sup>F100A</sup> | $5 \pm 1$ | $-7.7 \pm 0.1$ | $-17 \pm 5$ | $7 \pm 3$ | N/A |

| PKI<sub>5-24</sub> to ATPγN-saturated kinase | | | | | |
| --- | --- | --- | --- | --- | --- |
| | $K_d$ (μM) | $\Delta G$ (kcal/mol) | $\Delta H$ (kcal/mol) | $-T\Delta S$ (kcal/mol) | $\sigma$ |
| PKA-C<sup>WT</sup> | $0.16 \pm 0.02$ | $-9.33 \pm 0.07$ | $-13.9 \pm 0.5$ | $4.6 \pm 0.4$ | $106 \pm 18$ |
| PKA-C<sup>F100A</sup> | $2 \pm 1$ | $-7.9 \pm 0.3$ | $-17 \pm 1$ | $9 \pm 1$ | $3 \pm 1$ |

## Notes

### Competing Interest Statement

The authors have declared no competing interest.

### Summary of Updates

This revision includes revised figures 1 and 8 in the main text and a new supplementary figure 6. Also, the main text was revised for a broader audience and new citations were added.

